# Exposure to (*Z*)-3-hexenol primes tobacco plants for faster and stronger defense without negatively affecting their ability to grow and reproduce

**DOI:** 10.64898/2026.03.15.711883

**Authors:** Bipana Paudel Timilsena, Irmgard Seidl-Adams, Sarah Refi Hind, Gary Felton, James H. Tumlinson

## Abstract

Plants exposed to volatile signals from herbivore-infested neighbors can activate faster and stronger defenses against subsequent herbivore attack, a phenomenon called defense priming. However, the specific volatile components responsible for activating defense priming remain unclear. Here, we examined the role of green leaf volatiles (GLV) by silencing their biosynthesis using virus-induced gene silencing technique. Exposure to full blend of herbivore-induced plant volatiles (HIPV) primed receiver plants for enhanced production of all 5 groups of HIPV (GLV, monoterpenes, sesquiterpenes, aldoximes, and indole). When GLV production was silenced in emitter plants, receiver plants were no longer primed for terpene production. However, exposure to (*Z*)-3-hexenol (Z3HOL) alone primed receiver plants for terpene production. These results suggest that GLV are necessary, and Z3HOL alone is sufficient, to prime terpene production in receiver plants. Consistent with enhanced resistance, *Manduca sexta* larvae feeding on Z3HOL-or HIPV-primed plants consumed less leaf tissue and exhibited reduced growth compared with controls. Importantly, priming did not impose fitness costs, as Z3HOL-exposed plants showed normal growth but produced more seed capsules and seeds than control plants. Together, these findings suggest that Z3HOL alone is sufficient to prime plants for better defense without compromising their ability to grow and reproduce.

## 1. Introduction

Numerous studies have demonstrated that when plants are exposed to volatile signals from herbivore-infested neighbors, they tailor their defense reactions to incoming attack. After perceiving these volatile signals, some plants immediately exhibit direct induction of defense responses, such as upregulation of defense related genes and/or production of defense metabolites, even in the absence of herbivore attack (Farag & Paré 2002; Kost & Heil 2006). While others do not show immediate changes in defense status but respond more rapidly and more strongly following subsequent attack by herbivores, a phenomenon known as defense priming (Engelberth *et al*., 2004; Ton *et al*. 2007; Frost *et al*., 2008). The first evidence that plants can respond to volatile signals emitted by damaged neighbors was reported in the 1980s (Rhoades 1983; Baldwin & Schultz 1983). Since then, numerous studies have reported similar observations in both natural and laboratory settings (Heil & Karban 2010; Karban *et al*., 2014). However, after decades of research, it is still unknown what the distinguishing characteristics of the active compounds responsible for defense priming. Moreover, while most of the priming studies have focused on upregulation of defense genes and metabolites in primed plants, relatively little is known about how defense priming affects the performance of attacking herbivores and fitness of plants themselves.

Several different volatile compounds have been reported to induce defense priming in plants. The phytohormone ethylene was identified as the first gaseous signaling molecule in the early 1900s (Sasidharan 2022). Other volatile phytohormones, such as methyl jasmonate (meJA) and methyl salicylate (meSA), were also reported to induce defense responses in non-stressed neighboring plants (Farmer & Ryan 1990; Karban *et al*., 2000; Zavala & Baldwin 2004; Baldwin *et al*., 2006; Kessler *et al*., 2006; Tamogami *et al*., 2008). Yao *et al*. (2011) proposed involvement of additional signaling compounds beside volatile hormones (MeJA and MeSA) in plant defense priming. Accordingly, other candidate compounds for defense signaling include Green Leaf Volatiles (GLV), terpenes and aromatic compounds.

GLV (carbon-6 aldehydes, alcohols, and their esters) are among the most studied signaling molecules involved in defense priming (Cofer *et al*., 2018). Plants release GLV immediately after wounding or herbivory. Exposure to GLV activates several defense related genes and downstream metabolites (Zeringue 1992; Bate & Rothstein 1998; Engelberth *et al*., 2013). In addition to GLV, homoterpenes (*(E)-*3,8-dimethyl-1,4,7-nonatriene (DMNT) and (*E,E*)-4,8,12-trimethyl tridecane-1,3,7,11-tetraene (TMTT)) and monoterpenes (myrcene alone or blend of ocimenes consisting *(E)*-β-ocimene, *(Z)*-β-ocimene and allo-ocimene) have also been shown to induce defense related genes and meJA accumulation in exposed plants (Arimura *et al*. 2000; Godard *et al*., 2008; Meents *et al*. 2019). Similarly, exposure to physiologically relevant concentrations of indole (50 ng hr^-1^) also primes neighboring plants for enhanced production of JA, JA isoleucine (JA-Ile), abscisic acid (ABA) and plant volatiles following subsequent herbivore damage (Erb *et al*. 2015).

Yet most studies on priming, to date, have been performed using high doses of synthetic volatiles in airtight containers (Bate & Rothstein 1998; Engelberth *et al*. 2004). This high dose volatile exposure does not realistically represent what occurs in nature (von Mérey *et al*. 2011) and it also damages the plant tissue. The airtight container itself imposes physiological stress on plants. To address these limitations, a few studies attempted to mimic natural exposure by using genetically manipulated plants or slow-release dispensers but reported negative or unexpected results. For example, under field conditions, GLV-exposed plants experienced increased herbivore attraction and damage compared to unexposed control plants (von Mérey *et al*. 2011). Similarly, *N. attenuata* plants genetically impaired to emit GLV or terpenes failed to prime neighboring plants for defense gene expression and defense metabolite production following a single wounding event (Paschold *et al*., 2006). In contrast, volatiles from detached leaves of *Artemisia tridentata*, containing both GLV and terpenes, primed *N. attenuata* for enhanced proteinase activity only under continuous damage by *M. sexta* larvae (Kessler *et al*. 2006). These contrasting results suggest that, beyond volatile composition, the frequency and intensity of herbivore damage play a critical role in shaping plant defense and priming responses. In tobacco, continuous damage on plants built up JA production which is necessary to induce herbivore resistance mediated by both HIPV and defensive metabolites, such as nicotine and trypsin proteinase inhibitor (TPI) (Ziegler *et al*., 2001; Zavala *et al*., 2004; Zavala & Baldwin 2004; Halitschke *et al*., 2004). By contrast, a single wounding event elicits a quick burst of JA accumulation that rapidly returns to basal levels and is insufficient to confer herbivore resistance (Ziegler *et al*., 2001).

In a previous study, we found that the complete blend of HIPV primes neighboring *N. benthamiana* plants for enhanced JA and volatile production following subsequent herbivore challenge (Paudel Timilsena *et al*., 2020). Here we examined the role of GLV by silencing their biosynthetic pathway using virus-induced gene silencing (VIGS). We also exposed plants to individual GLV at physiologically relevant concentrations by using slow-release dispensers. In addition to quantifying defense metabolites, we also evaluated whether defense priming affects herbivore performance on primed plants, as well as the growth and reproduction of the plants themselves.

## 2. Materials and Methods

### 2.1 Plant materials and growing condition

Wild type (WT) tobacco plants, *N. benthamiana* (Solanaceae), were grown from seeds in individual pots (7.5 cm height, 9 cm diameter) filled with commercial potting soil (Pro-mix® PGX, Quakertown, PA) supplemented with Osmocote slow-release fertilizer (8-45-14 N-P-K, Scotts Company, Marysville, OH, USA). WT plants were grown in a greenhouse at 26 ± 2 °C under a 16 h light/ 8 h dark photoperiod. Plants used for experiments were 3-4 weeks old and had 4 fully developed leaves.

All VIGS plants were grown in a growth chamber (20 ± 2 °C and 16 h light/ 8 h dark) post VIGS inoculation. VIGS technique was used to silence the NbLOX2 gene as described in section below. Two days before the experiment, VIGS plants were transferred to and kept under the greenhouse conditions described above (26 ± 2^°^C, 16 h light/ 8 h dark). Transferring VIGS plants to the greenhouse condition with high temperature (26 ± 2°C) and high light intensity did not change the silenced phenotype (Nethra *et al*., 2006).

### 2.2 Insect and regurgitant

The Schilder lab at Penn State University, PA, generously provided *M. sexta* eggs. *M. sexta* neonates were reared on artificial diet under a 16 h light/ 8 h dark photoperiod at 25 ± 2°C. Oral secretions were collected from fourth- to fifth-instar *M. sexta* larvae feeding on *N. benthamiana* leaves for at least 24 h using previously described methods (Paudel Timilsena *et al*. 2020). Briefly, regurgitant was collected by squeezing larvae’s head and placing a pipette on their mouthpart. The collected regurgitant was immediately diluted with deionized water (1:1 v/v) and stored at -80 °C until use.

### 2.3 Chemicals

(*Z*)-3-hexenal (50%, SAFC, USA), (*Z*)-3-hexenol (>98%, Aldrich), (*Z*)-3-hexenyl acetate (>98%, Aldrich) were used to make slow-releasing dispensers. Hexane (>98.5%, J.T.Baker, USA) and dichloromethane (99.9%, OmniSolv, Germany) were used to elute volatiles from Super-Q (Alltech, USA) filters, and to wash Super-Q filters and glassware. Nonyl acetate (>97%, Aldrich, USA) was used as an internal standard to quantify volatiles. (*E)*-2-hexenal (98%, Bedoukian Research Inc., USA), linalool (97%, Aldrich), limonene (97%, Aldrich), (1, 8-) cineol (99%, Sigma-Aldrich), myrcene (90%, Aldrich), α-pinene (99%, Aldrich), β-pinene (99%, Aldrich), *(E)-*β-farnesene (99%, mixtures of isomers, Sigma-Aldrich) were used as reference standards for compounds identification based on mass spectra and retention time comparison using GC-MS and GC-FID, respectively.

### 2.4 Manipulation of HIPV release by emitter plants

We previously showed that exposure to the whole blend of HIPV primed neighboring plants for enhanced defense responses after subsequent challenge with herbivore regurgitant. To investigate the role of GLVs in priming, we generated emitter plants whose HIPV profile was lacking GLVs by silencing the lipoxygenase (LOX) gene in the hydroperoxide lyase pathway (HPL) branch of the oxylipin pathway (Bate & Rothstein 1998). We used virus-induced gene silencing (VIGS) to generate these GLV-compromised plants. These plants were exclusively used as emitter plants to produce defined volatile profiles at physiologically relevant concentrations. Therefore, any possible side effects of the VIGS would not affect their experimental role.

GLV are carbon-6 aldehyde, alcohol and their esters produced by the HPL branch of the oxylipin pathway. We silenced the homolog of the *N. attenuata* NaLOX2 gene in *N. benthamiana* plants (accession no. AY254348; Halitschke & Baldwin, 2003). Following herbivore damage, LOX2-silenced plants produced greatly reduced amounts of GLV, comparable to basal levels (**Fig. 2B**). Therefore, we used herbivore damaged LOX2 silenced plants as emitter plants and wild type (WT) as receiver plants. After 48 hours of exposure to volatiles, we measured the defense metabolites; volatiles and JA, in the WT receiver plants.

### 2.5 Plasmid construction for VIGS

Constructs used for this study include TRV::LOX2 (LOX2, lipoxygenase gene from *N. benthamiana*), TRV::GFP (GFP, green fluorescent protein, a non-plant gene obtained from jellyfish *Aequorea victoria* (Haseloff *et al*., 1997)) and TRV::PDS (PDS, phytoene desaturase from *N. benthamiana*). TRV::GFP was used as a VIGS control and TRV::PDS as a visual marker to confirm VIGS efficiency in all our VIGS experiments (Supplementary Fig. **S1**).

For cloning individual genes into pTRV2-LIC, the gene of interest (GOI) was amplified using the primers: 5’ CGACGACAAGACCCT (LIC vector adaptor) – GOI specific sequence-3’ and 5’-GAGGAGAAGAGCCCT (LIC vector adaptor) – GOI specific sequence - 3’ (Table **S1,** Dong *et al*., 2007). The insert size of each target gene was 300 bp in the TRV2-LIC vector. The PCR product of GOI was purified with a PCR clean-up kit. The 1000 ng of TRV2-LIC (pYY13) vector plasmid was digested by PstI-HF restriction enzyme in a 50 μL reaction, incubated for 1 h at 37 °C and purified with PCR-cleanup kit. A total of 50 ng of purified PCR product was treated with T4 DNA Polymerase in 1X reaction buffer containing 5 mM dATP and 5 mM Dithiothreitol (DTT) for 30 min at 22 °C followed by 20 min of inactivation of T4 DNA polymerase at 70 °C. A total of 500 ng of digested and purified TRV2-LIC (pYY13) plasmid was treated with T4 DNA polymerase and dTTP instead of dATP. Then the PCR product and TRV2-LIC (pYY13) vector were mixed in 1:1 ratio and incubated at 65 °C for 2 min and then 22 °C for 10 min. The mixture was then transformed into *Escherichia coli* strains TOP10 according to One Shot® Top10 Competent Cells protocol (Thermo Fisher Scientific). A total of 100 μL transformation reaction was spread onto LB plates containing 50 μg mL^-1^ of Kanamycin and incubated at 37 °C for up to 48 h. Transformants were selected by PCR and confirmed by running agarose gel electrophoresis. The plasmids from the confirmed transformants were isolated using Mini-Prep kit and were introduced into competent *Agrobacterium tumefaciens* strains GV2260 using the electroporation method by using a MicroPulser Electroporation according to the manufacturer’s (Bio-Rad) instructions.

### 2.6 Virus inoculation

Agrobacterium-mediated inoculation was conducted as described previously (Senthil-Kumar & Mysore, 2014). *Agrobacterium tumefaciens* strains GV2260 cells transformed with individual constructs were grown in LB (Luria-Bertani) broth, harvested and resuspended in MES (2-(N-morpholino) ethane-sulfonic acid), and incubation at room temperature for 3 h. Suspensions of Agrobacterium carrying pTRV1 and pTRV2 carrying GOI were mixed in 1:1 (v/v) ratio and infiltrated into leaves using a needleless syringe. Inoculated plants were grown in a growth chamber at 20 °C for 4 weeks.

### 2.7 Volatile dispenser

Plants were exposed to individual GLV for 48 hours using a slow-release dispenser designed to release physiologically relevant amounts for 5-7 days. Briefly, synthetic compound of interest was mixed with lanolin and 25 μL of the mixture was placed into a GC-insert. The GC-insert was immediately placed into the GC-vial and capped. A 1 μL, 64 mm micropipette was then inserted into the GC-vial. The emission rates were adjusted to physiologically relevant levels. For example, Z3HOL was released at 500 ng h^-1^ (mean, n = 6), which closely mimicked the emission rate observed in *M. sexta* damaged plant at 48 h post infestation (Supplementary Fig. **S2**). These calibrated dispensers were used as a source of volatile throughout the priming experiments.

### 2.8 Treatments on receiver plants, plant volatile collection and analysis

To examine the role of GLV in plant defense priming, WT receiver plants were exposed to volatiles emitted from *M. sexta* damaged WT or LOX2-silenced or GFP-silenced (VIGS control) plants as described in Paudel Timilsena *et al*. (2020). Briefly, two *N. benthamiana* plants were placed inside a bell jar (10 L); one was treated as an emitter plant and the other as a receiver plant (**Fig. 1**). Emitter plants were enclosed inside the wire case to prevent herbivore escape. In volatile exposure treatments, emitter plants were continuously damaged by *M. sexta* feeding, whereas control treatment, emitter plants remained undamaged. Volatiles emissions from undamaged plants were below the detection limit of GC-FID (Supplementary **Fig. S3**).

**Fig. 1.**
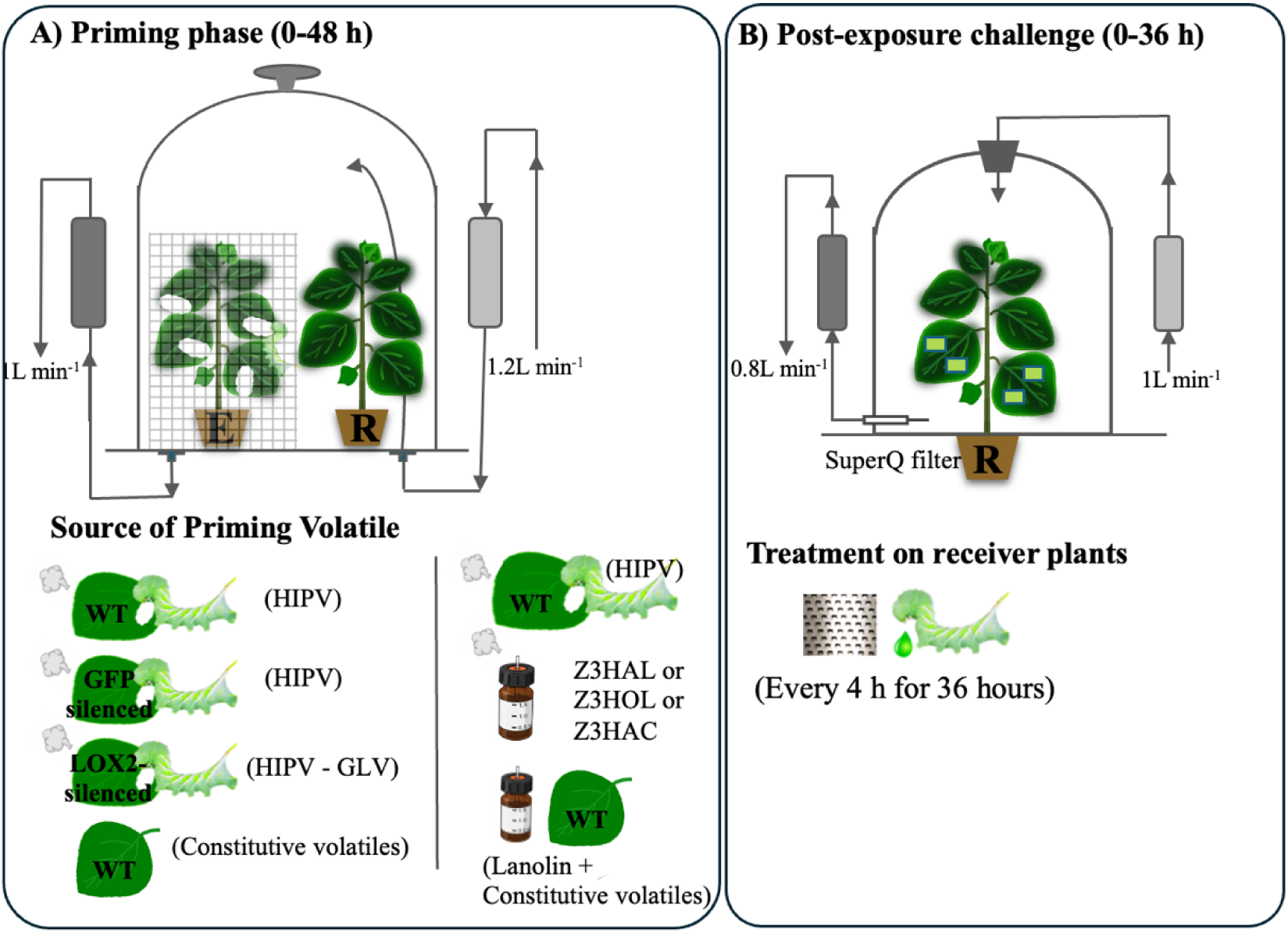
Experimental design for GLV-mediated defense priming. **(A) Priming phase (0-48 h):** In the first experiment, receiver plants (R) were exposed to volatiles from *M. sexta*-damaged WT LOX2-silenced, or GFP-silenced (VIGS-control) emitter plants (E). Receiver plants exposed to undamaged WT plants served as the negative control. In separate experiments, receiver plants were exposed to synthetic GLV [(*Z*)-3-hexenal (Z3HAL), (*Z*)-3hexenol (Z3HOL), or (*Z*)-3-hexenyl acetate (Z3HAC)] via slow-release dispensers. Receiver plants exposed to lanolin dispensers and undamaged WT plants as negative control, while those exposed to *M. sexta*-damaged WT served as positive control. (**B) Post-exposure challenge (0-36 h):** After priming, receiver plants (R) were transferred to clean bell jars and subjected to repetitive wounding every 4 h for 36 h using a cheese grater. Wounds were immediately treated with 10 μL of *M. sexta* regurgitant (1:1 v/v) to simulate herbivory. Volatiles were collected via SuperQ filters.

To examine the role of individual GLV, a slow-release dispenser loaded with the compound of interest was used as the volatile source and placed inside the bell jar with a receiver plant. Control plants were exposed to undamaged WT plant together with a dispenser containing lanolin only. To expose receiver plants to whole blend of volatiles, emitter plants were damaged by *M. sexta* as described above. During the priming phase, there was continuous airflow inside the bell jar (push air: 1.2 L min^-1^, pull air: 1 L min^-1^).

After 48 h of exposure to volatiles, receiver plants were transferred to individual clean bell jars (8 L) and repeatedly wounded every 4 h during the light period for two consecutive days (Paudel Timilsena *et al*. 2020). During each wounding event, two leaves were damaged once on each side of the mid rib using a fine cheese grater (3 cm × 2 cm) that made ∼180 holes per wounding site. Thus made wounds were immediately treated with 10 μL of *M. sexta* regurgitant (diluted with water, 1:1 v/v), which was identified previously to mimic insect herbivory (Heil *et al*. 2001, 2012).

### 2.9 Headspace volatiles collection and analysis

Following simulated herbivory, headspace volatiles from individual receiver plants were collected at 4, 8, 12, 16, 20, 24, 28, 32 and 36 h using a dynamic headspace sampling system as previously described by Paudel Timilsena *et al*. (2020). Briefly, clean, filtered air was pushed into the bell jar at 1 L min^-1^, and outgoing air was pulled through glass filter containing 30 mg of Super-Q adsorbent (Alltech, Deerfield, IL, USA) at 0.8 L min^-1^. Trapped volatiles on the filters were eluded using 120 μL of a hexane: dichloromethane (1:1 v/v) containing nonyl acetate (4 ng μL^-1^) as an internal standard. Volatile compounds were quantified using gas chromatograph coupled with a flame ionization detector (Agilent 6890 GC-FID, Agilent, CA, USA) fitted with a capillary column (HP-5, 30 m × 0.25 mm ID × 0.25 μm film thickness; Supelco, PA, USA). Samples (1 μL) were injected using an automated injector. The GC oven was programmed from 40 °C (2 min hold) to 150 °C at 4 °C min ¹, then to 290 °C at 40 °C min ¹, with a final 3 min hold. Helium was used as the carrier gas at 14.79 psi. Peak areas were integrated using GC ChemStation software, and volatile quantities were calculated relative to an internal standard.

Compound identities were verified by GC-MS (Agilent 6890N GC coupled to a 5973N MS) using the same column and temperature program. (*Z*)-3-hexenal, (*Z*)*-*3-hexenol, (*Z*)*-*3-hexenyl acetate, (*E*)*-*2-hexenal, linalool, limonene, (1, 8-) cineol, myrcene, α-pinene, β-pinene, sabinene, and (*E*)*-*β-farnesene were verified by comparing retention times and mass spectra with synthetic standards. Remaining compounds were identified by matching spectra to the NIST library (GCMS Solution, Shimadzu, MD, USA).

### 2.10 Phytohormone JA extraction and quantification

We measured JA accumulation at the damaged site of receiver plants as an indicator of primed defense response. After 48 h of exposure to volatiles, receiver plants were damaged with simulated herbivory. The third leaf from the base was selected and mechanically wounded on both sides of the mid rib and immediately treated with *M. sexta* regurgitant. The damaged portion of the leaf was harvested 30 min after challenge. The tissue was flash frozen into liquid nitrogen and stored at -80 °C until analyzed. All the samples were ground into fine powder with pre-cooled mortar and pestle under liquid nitrogen. The powdered sample (100 mg) was transferred into a 2-mL screw-cap Fast Prep Tubes (Qbiogene, CA, USA) previously loaded with 1 g of Zirmil beads (1.1 mm; SEPR Ceramic Beads and Powders, NJ, USA), 400 μL extraction solvent (isopropanol: H2O, 2:1 v/v, adjusted to pH 3 with hydrochloric acid) and 20 μL dihydro-JA (10 ng μL^-1^) an internal standard and vortexed thoroughly. Then 1 mL of dichloromethane was added to each sample vortexed vigorously for 20 sec using a FastPrep FP 120 tissue homogenizer (Qbiogene) and centrifuged at 10,000 rpm for 2 min. The bottom organic phase (dichloromethane and isopropanol layer) was transferred to a 4-mL screw cap glass vial, dried under nitrogen gas. The dried extract was re-suspended in 200 μL diethyl ether: methanol (9:1 v/v) and then esterified by adding 3 μL trimethylsilyldiazomethane solution (2 M in hexanes, Andrich). The vial was then capped, vortexed, and incubated at room temperature for 30 min. The excess of trimethylsilyldiazomethane was inactivated by adding 3 μL of 2 M acetic acid. The vial was again capped, vortexed, and incubated at room temperature for 30 min. To collect methylated and volatilized JA, a SuperQ filter attached to the vacuum source was inserted into the vial through a cut in the septum. The vial was heated in a heating block to 180 °C for 2 min. The methylated and volatilized JA was then trapped on the SuperQ filter for 2 min at 400 mL min^-1^. Trapped JA was eluded with 150 μL of dichloromethane and analyzed using a gas chromatograph coupled to a mass spectrometer as previously described (Paudel Timilsena *et al*. 2020). The methylated product of endogenous JA was quantified by correlating the peak area of the compound with that of the dihydro-JA (internal standard) using GC Chemstation software.

### 2.11 Herbivore performance bioassay

First, we investigated the mortality rate of freshly hatched *M. sexta* on the plants that were exposed to Z3HOL and the volatiles from undamaged emitter plants (here after air-exposed). After 48 h of exposure, all the receiver plants were treated with 20 neonates per plant. The mortality rate of the neonates was recorded at 24 h after the insect treatment.

To analyze the rate of feeding consumption, two third-instar larvae of *M. sexta* were placed in an oblong container (30 cm × 15 cm) with four leaves of HIPV-exposed, Z3HOL-exposed or air-exposed plants. The containers were previously coated with a thin layer of 1% agar to prevent leaf wilting and lids had nylon mesh-covered holes for ventilation. Twenty-four h after the start of the experiment, the area of leaf tissue removed was analyzed using the software ImageJ (Schneider *et al*., 2012).

In a separate experiment, growth of third-instar larvae on Z3HOL-exposed or air-exposed plants was analyzed. After 48 h of exposure to volatiles, receiver plants were treated with third-instar larvae of *M. sexta* (4 plant^-1^) for 7 days and the insect weight gain was recorded at the beginning (day 0) and end of the experiment (day 7).

### 2.12 Herbivore behavioral response and dual choice bioassay

We characterized behavioral responses of *M. sexta* larvae to HIPV-exposed, Z3HOL-exposed and air-exposed control plants using a dual-choice assay in an oblong container (30 cm × 15 cm). The 3^rd^ and 4^th^ leaves from HIPV- or Z3HOL-exposed or unexposed tobacco plants (a combination of two treatments) were collected and placed on opposite sides of the container. The container previously was coated with a thin layer of 1% agar to prevent leaf wilting or drying out. Four 3^rd^ instar *M. sexta* larvae were released at the center of the container and covered with the lid containing nylon mesh for ventilation. Larvae were free to move inside the container and choose between two leaf sources. The number of *M. sexta* on each side of the container was recorded at 1, 6, 12 and 24 h.

### 2.13 Plant growth and development experiment

To assess the effect of volatile exposure on plant growth and development, growth parameters (such as plant height, number of secondary branches, secondary branch length) and reproduction parameters (such as number of seed capsules, seed mass and seed germination percentage) were measured. Two weeks after transplanting, *N. benthamiana* plants were exposed to Z3HOL using the slow release volatile dispenser. Dispensers were calibrated to release Z3HOL at the rate of 500 ng hr^-1^ for 1 week and were replaced weekly. Once the plants started to flower, volatile dispensers were removed. Control treatments were exposed to identical dispensers containing only lanolin. All the plants were kept in a greenhouse at 26 ± 2^°^C under 16L: 8D h photoperiod.

### 2.14 Statistical analysis

The volatile data from the time course experiments were analyzed using the mixed-model for repeated measures, SAS PROC MIXED (Saxton 1998), procedure in SAS statistical software (SAS Institute Inc. 2013; SAS/STAT 9.4 User’s Guide, NC, USA) as described previously (Seidl-Adams *et al*. 2015; Paudel Timilsena *et al*. 2020). For repeated measures analysis with Wald F-statistics, the best covariance structure was selected based on the Akaike Information Criterion (AIC) and Bayesian Information Criterion (BIC), and the degrees of freedom were adjusted using the Kenward-Roger method (Gomez *et al*. 2005). Since volatile emission follows diurnal rhythm (Rim *et al*. 2019), volatile data were divided into three subsets (Day-1, Night-1, and Day-2) and analyzed separately. For each dataset, two separate hypotheses were tested: 1) mean volatile emissions at particular time points were same for all treatments, 2) mean volatile emission for particular treatments were same at all time points. Then, the comparisons of interest were extracted and evaluated for significance using the Bonferroni adjustment for multiple comparisons.

All the other statistical analyses were conducted in R (R Core Team 2025). Data from insect feeding assays were analyzed using one-way ANOVA, after confirming normally distributed residuals (Shapiro-Wilk test), and homogeneity of variances (Levene’s F-test). When significant differences (P < 0.05) were found, pairwise post hoc comparisons of least squares means were conducted using Least Significant Difference (LSD) at P ≤ 0.05. Plant growth and development data were analyzed by Student’s *t*-tests or Welch two sample *t*-tests, depending on whether samples violated the assumption of equal variance or not. The nature of dual-choice assay with *M. sexta* larvae per case generated discrete data. Therefore, we used the nonparametric statistical procedure, Wilcoxon matched-pair test, to determine the difference in the number of larvae settled on either treatment.

## 3. Results

### 3.1 LOX2-silenced plant produces all the other volatiles except GLVs

Approximately 25 d after inoculation of early rosette-stage *N. benthamiana* with *Agrobacterium tumefaciens* GV3101 harboring pTRV::LOX2 (gene of interest), pTRV::GFP (vector control) and pTRV::PDS plasmid, new growth of PDS-silenced *N. benthamiana* plants became completely bleached, demonstrating successful gene silencing (Supplementary Figure **1)**. Silencing of NbLOX2 did not markedly affect plant growth compared to the vector control (NbGFP-silenced) or WT plants grown under the same environment conditions. To determine the silencing efficiency of NaLOX2, we analyzed volatiles emitted from *M. sexta* feeding damaged NaLOX2-silenced plants and compared them with similarly treated NaGFP-silenced plants (vector control) and WT plants. Headspace collection followed by GC-FID analysis shows no significance difference in mono- and sesquiterpene production by *M. sexta* fed WT, GFP-silenced and LOX2-silenced plants (Fig. **2A**). However, LOX2-silenced plants emitted only basal amounts of GLVs after *M. sexta* feeding, whereas WT and GFP-silenced plants showed robust GLV induction (Fig. **2A**).

**Fig. 2.**
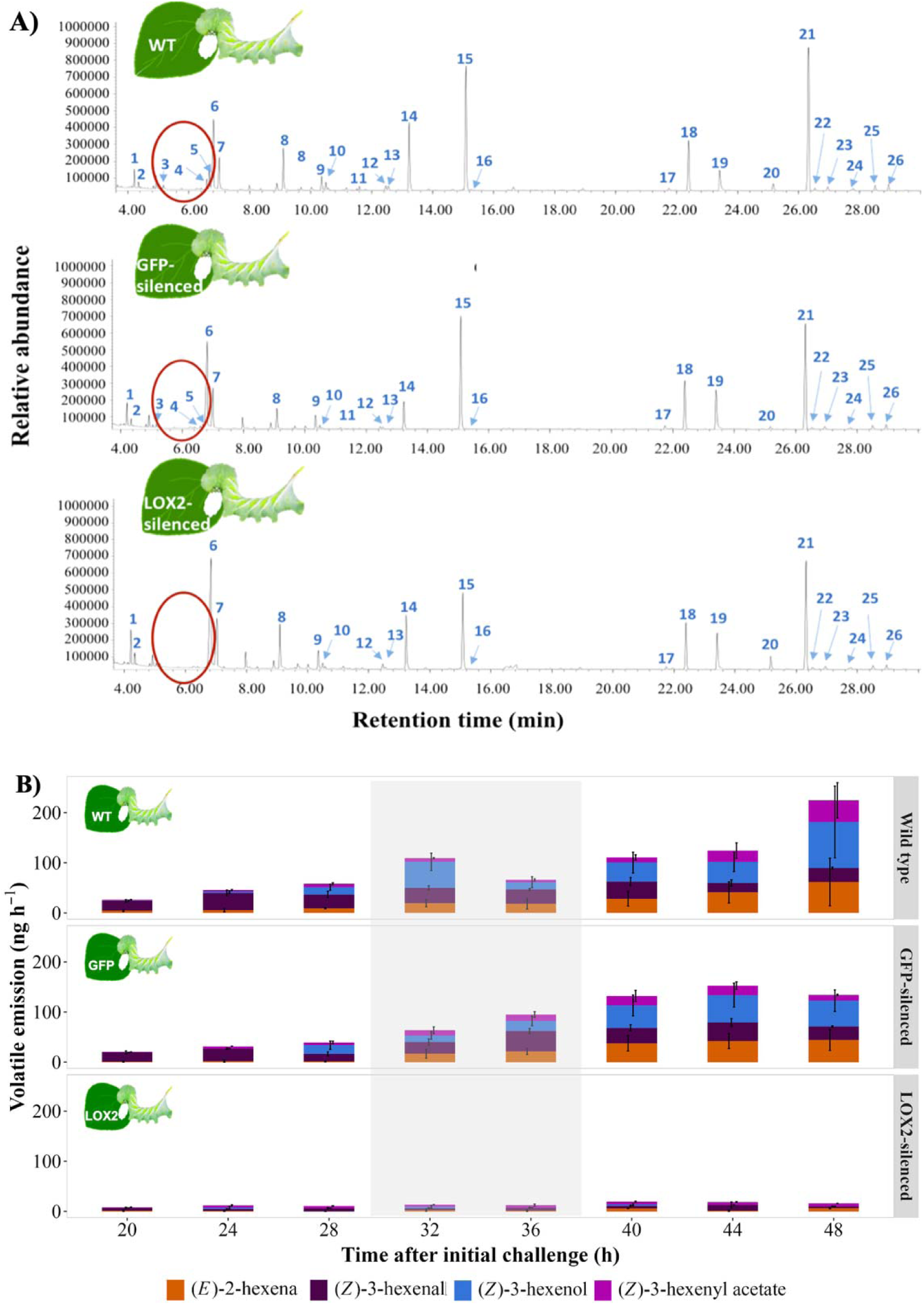
HIPVs emitted from *M. sexta* damaged *N. benthamiana* plants. **A)** Chromatographic profiles of HIPVs emitted by *M. sexta* damaged wild type (top), GFP-silenced (middle), and LOX2-silenced (bottom) plants. *N. benthamiana* plants were continuously damaged by fourth-instar *M. sexta* feeding for 48 h. The chromatogram represents HIPV profile collected from 24 to 28 h after feeding initiation. Five major families of HIPVs were induced by *M. sexta* feeding on *N. benthamiana*: **green leafy volatiles (GLVs):** (*Z*)-3-hexenal [3], (*E*)-2-hexenal [4], (*Z*)-3-hexenol [5], (*Z*)-3-hexenyl acetate [11], **monoterpenes:** α-pinene [8], sabinene [9], β-pinene [10], limonene [12], 1,8-cineole [13], (*E*)-β-ocimene [14], linalool [15], nonanal [16], **sesquiterpenes:** (*E*)-α-bergamotene [21], unknown sesquiterpene#1 [22], (*E*)-β-farnesene [23], aristolochene (4,4-di epi) [24], β-bisabolene [25], β-sesquiphellandrene [26], **aldoximes:** propyl aldoxime, 2 methyl-,syn-[1], propyl aldoxime, 2 methyl-,anti-[2], butyl aldoxime, 2 methyl-,syn- [6], butyl aldoxime, 2 methyl-,anti-[7], and **aromatic compound**: indole [17]. **A) Levels of GLVs emitted by *M. sexta* damaged plants.** Early fourth-instar *M. sexta* (one larva per plant) was feeding continuously on the plant for 48 hours. HIPVs collection started on 2nd day, 16 h after feeding initiation and continued up to 48 h in 4-h intervals. Values represent mean ± SE (n = 4). Shaded areas indicate nighttime volatile collections.

The main purpose of silencing GLV production was to investigate the role of GLV in defense priming. In previous priming experiments, receiver plants were exposed to HIPVs for 48 h (Paudel Timilsena *et al*. 2020). Therefore, we used the same time window of 48 h and confirmed that NbLOX2-silenced plants produce basal level of GLV after *M. sexta* feeding for 48 h (Fig. **2B**).

### 3.2 GLVs are necessary for terpene priming

WT receiver plants were exposed for 48 h to volatiles emitted from *M. sexta* damaged WT, LOX-2 silenced, or GFP-silenced plants or from undamaged WT plants. Following volatile exposure, all receiver plants were challenged (mechanical damage followed by *M. sexta* regurgitant application) and emitted volatiles were collected for 36 h at 4-h intervals. When GLV production was silenced in emitter plants, receiver plants produced significantly lower amount of mono- and sesquiterpenes compared with plants exposed to the whole blend of volatile (i.e, volatiles from *M. sexta* fed WT and GFP-silenced emitters). Receiver plants exposed to LOX2-silenced plants and the undamaged WT plants produced similar amounts of mono- and sesquiterpenes after challenge (Fig. **3A-B**).

**Fig. 3.**
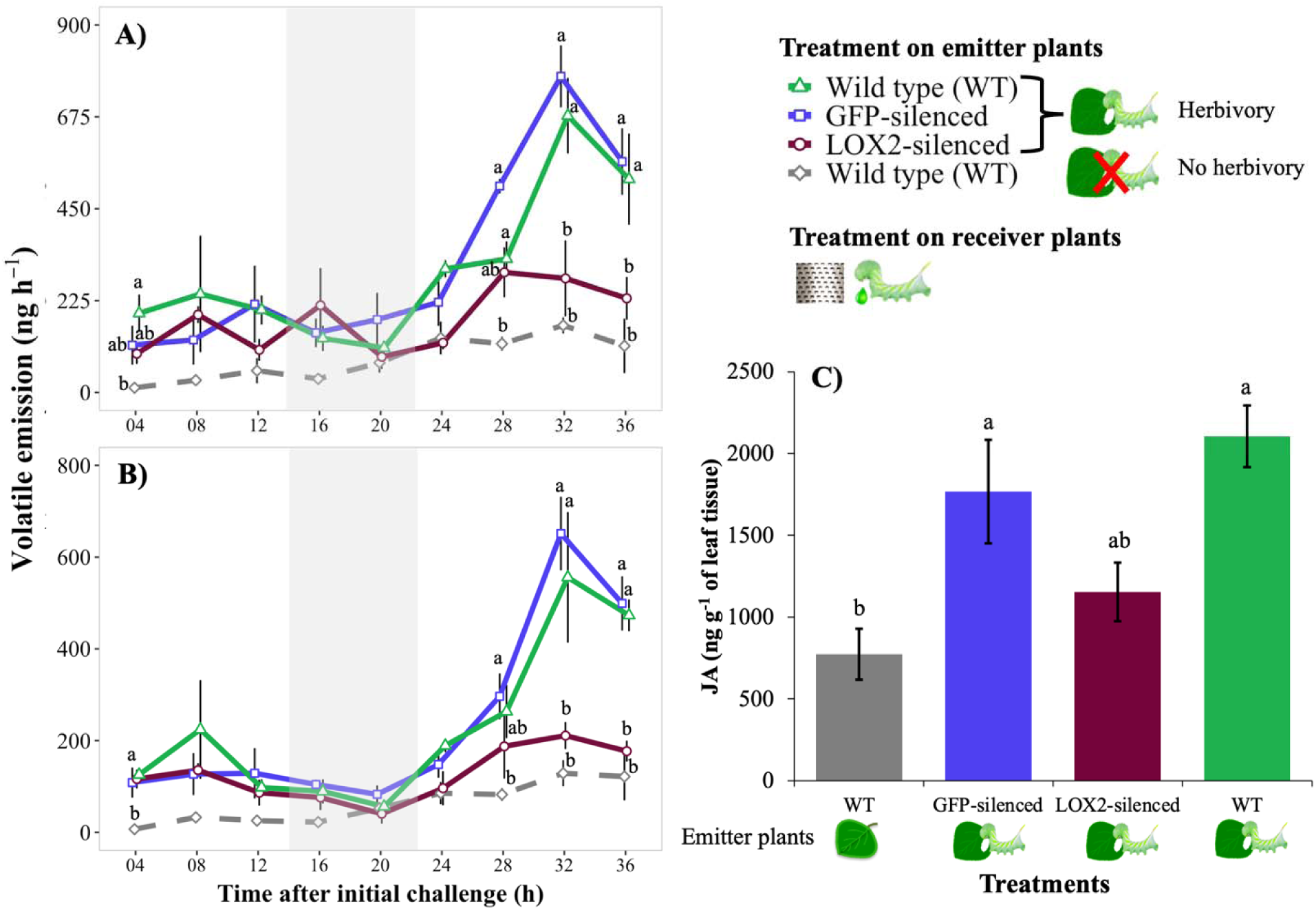
Receiver *N. benthamiana* plants did not get primed for terpene emissions when GLV emission was silenced in emitter plants. *N. benthamiana* plants were exposed to volatiles from *M. sexta* damaged WT (green), LOX2-silenced (dark red), GFP-silenced (blue) or undamaged (gray) plants for 48 h. After 48 h of exposure to volatiles, the receiver plants were transferred to individual clean bell jars and challenged repeatedly by mechanical wounding followed by *M. sexta* regurgitant application. The graph shows the total amount of **A)** monoterpenes and **B)** sesquiterpenes emitted from the challenged receiver plants at different time intervals after initial challenge. Values represent mean ± SE (n = 3). Data were analyzed with a mixed model for repeated measures. Different letters indicate significant (*p* < 0.05) differences between treatment within time points, with Bonferroni’s correction for multiple comparisons. Shaded areas indicate nighttime volatile collections. **C)** For JA analysis, after 48 h of exposure to volatiles, receiver plants were challenged with mechanical damage followed by *M. sexta* regurgitant application. Wounded portions of the leaf were harvested 30 min after challenge and JA accumulation in the harvested tissue was analyzed (ANOVA, Tukey HSD, n = 3, *p* < 0.01).

Similarly, JA accumulation at the damaged site of the receiver plants was only strongly primed after exposure to the whole blend of HIPV. Receiver plants exposed to *M. sexta* damaged WT and GFP-silenced emitters produced significantly higher amounts of JA than those exposed to undamaged emitter plants (Fig. **3C**). Although not statistically significant, receiver plants exposed to *M. sexta* damaged LOX2-silenced emitters produced 82.39 % and 53.11 % less JA than plants exposed to similarly treated WT and GFP-silenced emitters, respectively. Thus, removing GLV from the HIPV blend diminished priming but did not completely abolish JA induction, likely because other HIPV components still contribute to the response.

Together, these results suggest that GLV is necessary for full priming of mono- and sesquiterpene production and contribute subsequently to JA accumulation in neighboring *N. benthamiana* plants (Fig. **3**).

### 3.3 Exposure to (*Z*)-3-hexenol alone primes terpene emission

*M. sexta* damaged *N. benthamiana* plants emit four major GLV: (*Z*)-3-hexenal, (*E*)-2-hexenal, (*Z*)-3-hexenol and (*Z*)-3-hexenyl acetate (Supplementary Fig. **S2D**). To determine whether any individual GLV is sufficient for priming, plants were exposed to the physiologically relevant concentration of each GLV using slow-release dispensers.

To test whether (*Z*)-3-hexenol (here after Z3HOL) acts as a priming agent between plants, healthy *N. benthamiana* plants were exposed to synthetic Z3HOL at the release rate of 500 ng hr^-1^ for 48 h. Following exposure, receiver plants were repeatedly challenged (mechanical damage followed by *M. sexta* regurgitant application) and volatiles were collected for 36 h. On the second day of challenge, emission of sesquiterpene was significantly enhanced in Z3HOL-exposed plants, demonstrating that exposure to synthetic Z3HOL alone is sufficient to prime mono- and sesquiterpene emission in *N. benthamiana* plants (Fig. **4A-B**). However, exposure to the full blend of volatiles resulted in stronger terpene than exposure to Z3HOL alone across all sampling periods. These results suggest that while Z3HOL can independently induce priming, primed responses are amplified when the plants are exposed to the complete blend of volatiles. Although (*Z*)-3-hexenal and (*Z*)-3-hexenyl acetate were also tested in separate experiments, Z3HOL was the only compound tested that primed *N. benthamiana* plants for induced volatile emissions and JA production (**Fig. S4-5**).

**Fig. 4.**
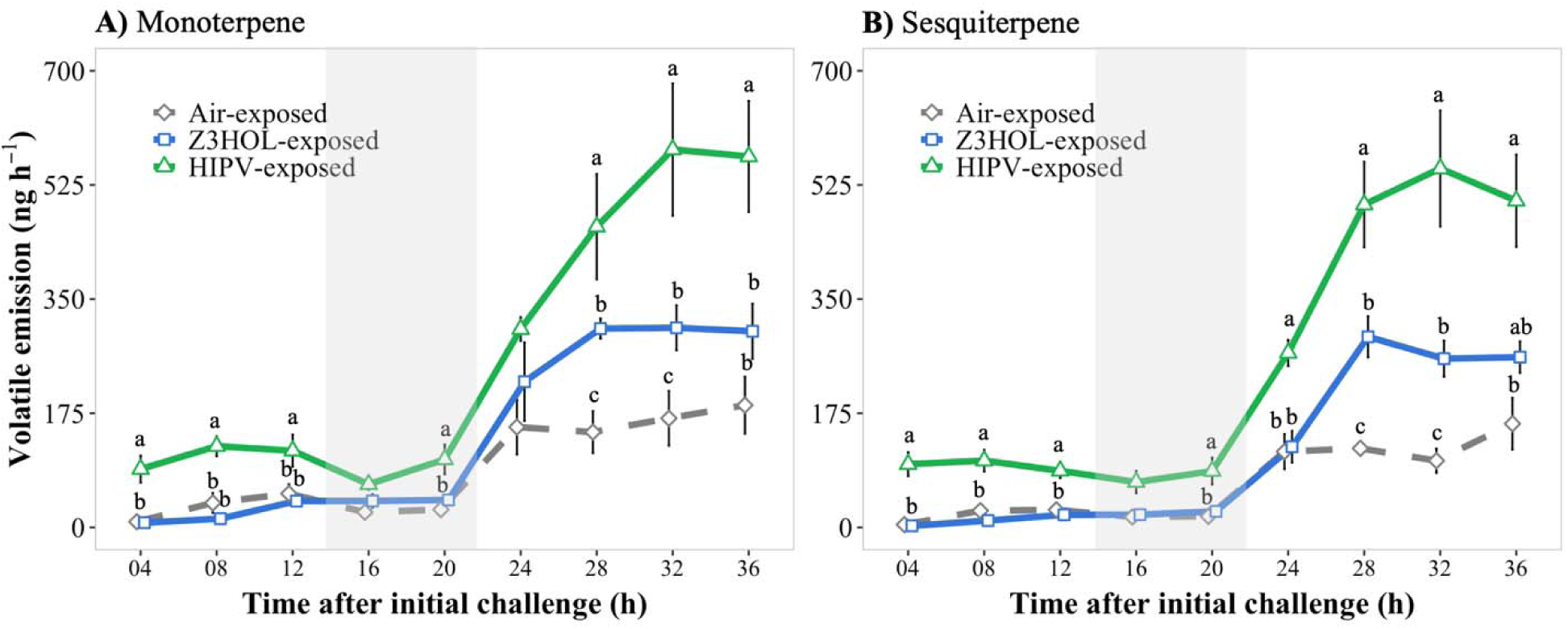
Exposure to Z3HOL alone primed *N. benthamiana* plants for enhanced production of mono and sesquiterpenes. *N. benthamiana* plants were exposed to volatiles from *M. sexta* damaged WT (green), undamaged WT plants (gray) or synthetic Z3HOL (red) for 48 h. After 48 h of exposure to volatiles, the receiver plants were transferred to individual clean bell jars and challenged repeatedly by mechanical wounding followed by *M. sexta* regurgitant application. The graph shows the total amount of **A)** monoterpenes and **B)** sesquiterpenes emitted from the challenged receiver plants at different time intervals after initial challenge. Values represent mean ± SE (n = 4). Data were analyzed with a mixed model for repeated measures. Different letters indicate significant (*p* < 0.05) differences between treatments within time points, with Bonferroni’s correction for multiple comparisons. Shaded areas indicate nighttime volatile collections.

### 3.4 *M. sexta* larvae performed poorly on volatile exposed plants

To further examine whether Z3HOL alone is sufficient to prime defense responses against herbivory, *N. benthamiana* plants were exposed for 48 h to this volatile at comparable amounts that were emitted from *M. sexta* damaged plants, i.e 500 ng hr^-1^. After exposure, plants were infested with freshly hatched *M. sexta* neonates (15 per plant), and neonate mortality was recorded 48 h after release. Neonate mortality rate did not differ significantly among treatments (F = 3.374, *p* = 0.054, n = 8; **Fig. 5A**). However, the overall neonate mortality rate was very high, above 50%, across all treatments. We therefore evaluated *M. sexta* performance using early third-instar larva. Larvae were fed on tobacco leaves for at least for 24 h prior to the experiment, and only healthy, actively feeding larvae were selected. The mortality rate of these larvae was lower than that of freshly hatched neonates (Supplementary Fig. **S6**).

**Fig. 5.**
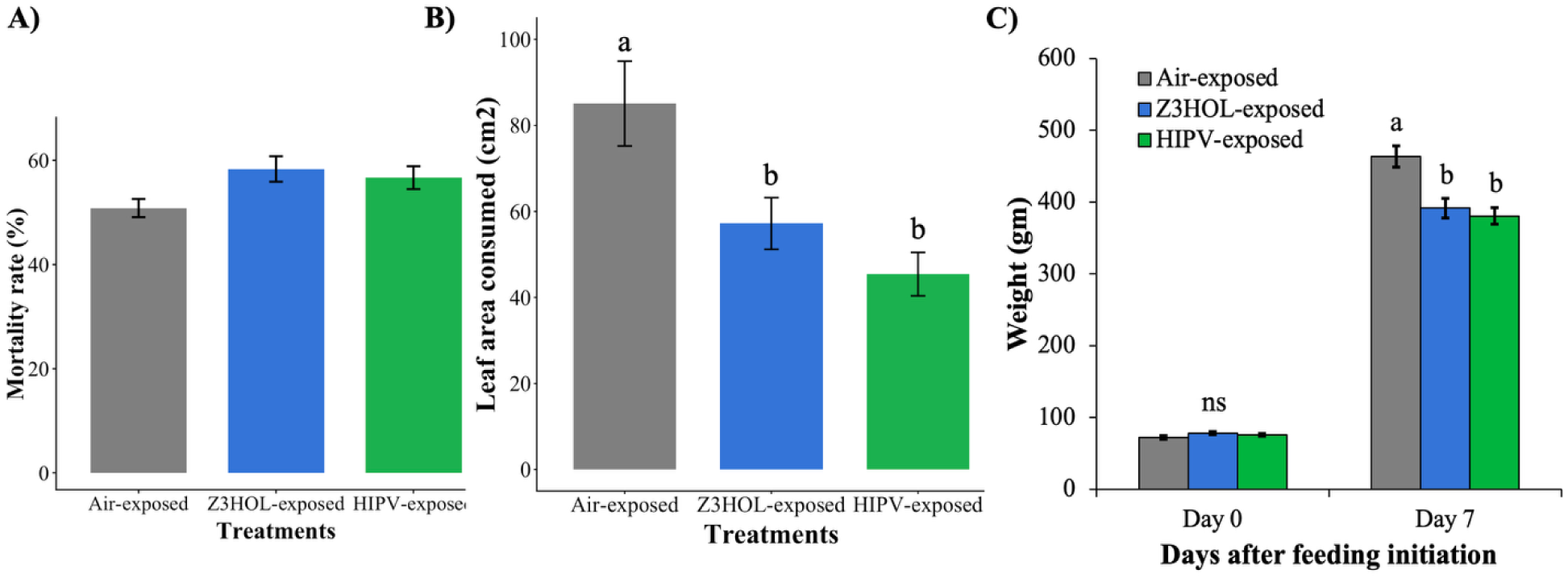
Volatile (Z3HOL and HIPV) exposure primed defenses against *M. sexta* larvae. *N. benthamiana* plants were exposed for 48 h to volatile from undamaged plant (air-exposed, gray), 500ng of Z3HOl (Z3HOL-exposed, blue) or HIPVs from *M. sexta* damaged plants (HIPV-exposed, green). **A)** Mortality rate of freshly hatched *M. sexta* neonates 48 h after the neonate release on receiver plants (ANOVA, F = 3.374, n = 8, *p* = 0.054), **B)** Amount of leaf area consumed by two early-3rd-instar *M. sexta* larvae during 24 h feeding period (ANOVA, Tukey HSD, F = 11.27, n = 6, *p* < 0.001), **C)** Larval weight gain at 0 and 7 days after feeding on receiver plants (ANOVA, Tukey HSD, F = 7.84, n = 6, *p* < 0.01). The bar represents the mean ± SE. Letters indicate significant differences among different treatments.

*N. benthamiana* plants exposed to volatiles (either Z3HOL or HIPV) showed enhanced defense against freshly molted 3^rd^-instar *M. sexta* larvae, as indicated by significantly reduced leaf consumption and larval weight gain. *M. sexta* larvae feeding on Z3HOL- and HIPV-exposed plants gained significantly less weight than those feeding on air-exposed control plants, with no significant difference between the Z3HOL and HIPV treatments (F = 11.27, *p* < 0.001, Fig. **5b**). Similarly, during a 24 h no-choice feeding assay, *M. sexta* consumed significantly less leaf materials when they were placed on detached leaves of Z3HOL- and HIPV-exposed plants compared to leaves from air-exposed plants (F = 7.84, *p* = 0.004; Fig. **5C**). Leaf consumption was decreased by 46.65% and 32.78% on HIPV- and Z3HOL-exposed plants, respectively, compared to air-exposed controls.

### 3.5 *M. sexta* larvae prefer air-exposed plants over volatile-exposed plants

We characterized behavioral responses of *M. sexta* larvae to air-exposed, Z3HOL-exposed or HIPV-exposed *N. benthamiana* plants using dual-choice assays with all the possible combinations. At the 1 and 6 h after assay initiation, the 3^rd^-instar *M. sexta* larvae did not discriminate between leaves from differently treated plants in the dual choice assay. However, at 12 and 24 h, significantly higher numbers of larvae settled on leaves from air-exposed plants, suggesting reduced preference to Z3HOL- or HIPV-exposed plants **(Fig. 6A)**. Similarly, when larvae were given a choice between Z3HOL- and HIPV-exposed plants, significantly more larvae settled on Z3HOL-exposed plants than HIPV-exposed plant at 6, 12 and 24 h **(Fig. 6A),** suggesting that HIPV-exposure might have primed additional changes or stronger responses in the receiver plants that further reduced larval preference.

**Fig. 6.**
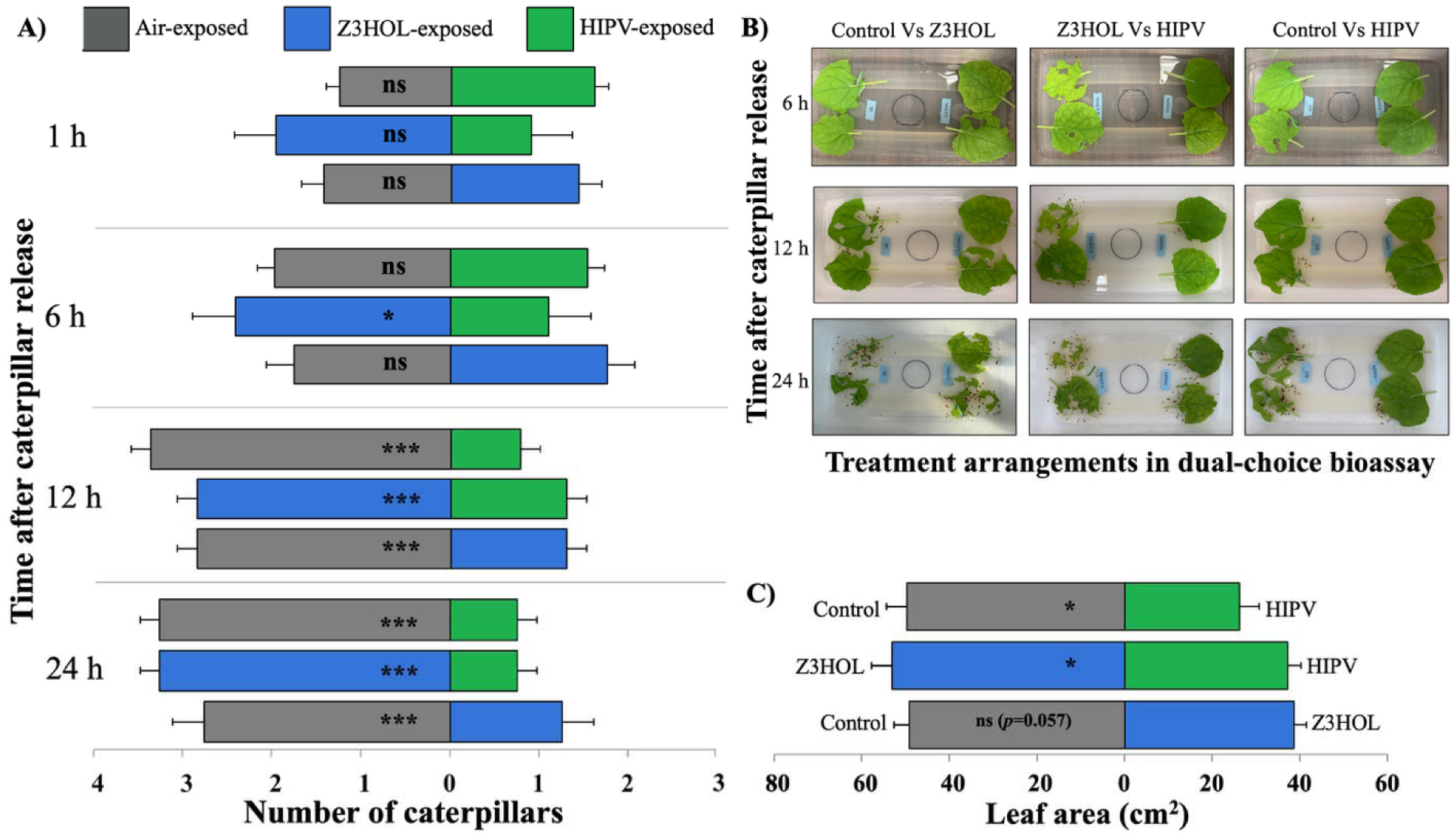
Feeding preference of *M. sexta* larvae is influenced by volatile exposure treatments. **A)** The preference of *M. sexta* larvae on detached leaves of air-exposed, Z3HOL-exposed or HIPV-exposed plants. Four early-3^rd^-instar larvae were used for each replication, with 8 replications in total. The bar represents mean ± SE (n = 8). * and *** indicate significant difference between treatments at *p* < 0.05, *p* < 0.001, respectively, according to Wilcoxon rank sum test. **B)** Representative differences in amount of leaf material consumed by four 3^rd^-instar *M. sexta* larvae. Pictures were taken at 6, 12 and 24 h after the release of caterpillars in the tub. **C)** Leaf area consumed by four 3^rd^-instar *M. sexta* larvae during 24 h feeding period (dual-choice bioassay). The bar represents mean ± SE (n = 8). * Indicates significant differences between treatments at *p* < 0.05 according to Student’s *t* test.

In addition to larval settlement, volatile exposure also significantly influenced the amount of leaf consumption. During a 24 h feeding period, *M. sexta* larvae consumed significantly lower amounts of leaf materials on HIPV-exposed plants when given a choice between air-exposed vs HIPV-exposed plants, as well as between Z3HOL-exposed vs HIPV-exposed plants **(Fig. 6B-C)**. When given a choice between air-exposed and Z3HOL-exposed plants, *M. sexta* larvae consumed 12% less leaf materials from Z3HOL-exposed plants, although this difference was not statistically significant (*p* = 0.057). Together, these results indicate that both HIPV- and Z3HOL-exposed plants are less preferred by *M. sexta* larvae compared to air-exposed plants, with HIPV-exposure eliciting the strongest deterrent effect.

### 3.6 *N. benthamiana* plants exposed to Z3HOL or HIPV exhibit enhanced defense responses

To understand the reduced performance and preference of *M. sexta* larvae on Z3HOL-and HIPV-exposed plants, we quantified the levels of the key defense phytohormone, jasmonic acid (JA), in exposed and air-exposed plants before and after herbivore damage. Herbivore damage induced accumulation of JA in all plants at the wound site but the magnitude of induction was significantly greater in Z3HOL- and HIPV-exposed plants. At 30 minutes after challenge (mechanical damage + *M. sexta* regurgitant application), JA levels were 2.57 and 2.63 folds higher in Z3HOL- and HIPV-exposed plants, respectively, compared with air-exposed plants (Fig. **7**).

**Fig. 7.**
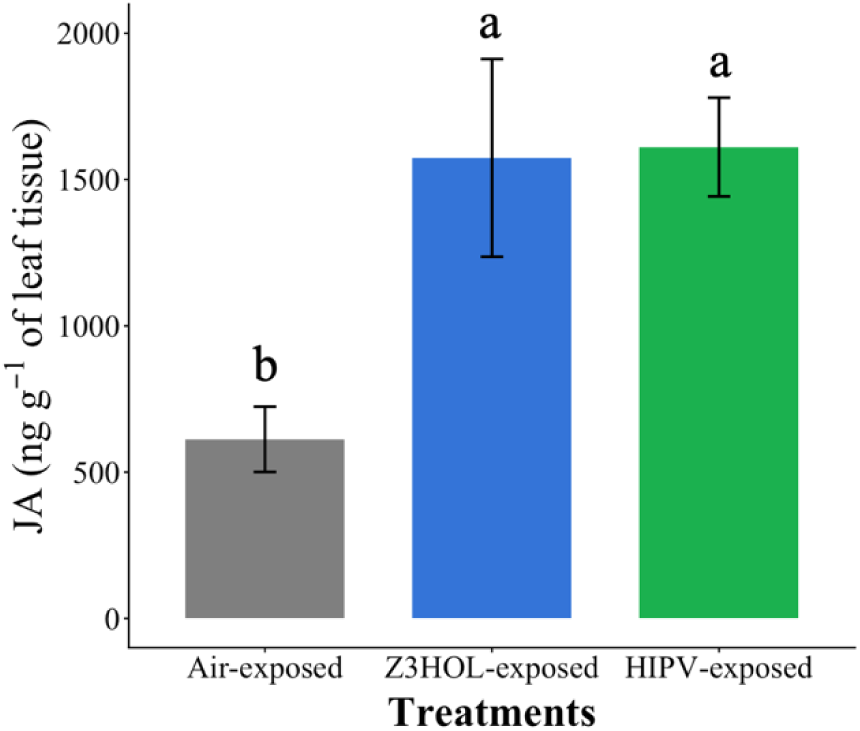
Volatile (Z3HOL or HIPV) exposed plants respond more strongly to simulated herbivory than air-exposed plants. *N. benthamiana* plants were exposed for 48 h to volatile from undamaged plant (air-exposed, gray), 500 ng of Z3HOL (Z3HOL-exposed, blue) or HIPVs from *M. sexta* damaged plants (HIPV-exposed, green). After 48 h of exposure to volatiles, receiver plants were challenged with mechanical damage followed by *M. sexta* regurgitant application. Wounded portions of the leaf were harvested 30 min after challenge for JA analysis (ANOVA, Tukey HSD, n = 4, *p* < 0.05). The bar represents the mean ± SE. Letters indicate significant differences among different treatments.

### 3.7 (*Z*)-3-hexenol exposure enhanced the reproductive fitness of the plant

To explore whether the exposure to Z3HOL affects plant growth and development, tobacco plants were exposed to Z3HOL from two-week after transplanting until the onset of flowering under insect-free conditions. The Z3HOL dispensers were replaced weekly to maintain a consistent supply of Z3HOL. Z3HOL exposure did not affect vegetative growth, as exposed plants showed no significant differences in plant height, number of secondary branches per plant, and average length of secondary branch compared with air-exposed control plants (Figure **8A-C**). In contrast, reproductive performance was significantly enhanced by Z3HOL exposure. At harvest, Z3HOL-exposed plants produced significantly more seed capsules and greater seed mass compared to unexposed control plants. Specifically, Z3HOL-exposed plants had about 13% more seed capsules and a 24% increase in seed yield compared to control plants (Figure **8E-F**).

**Fig. 8.**
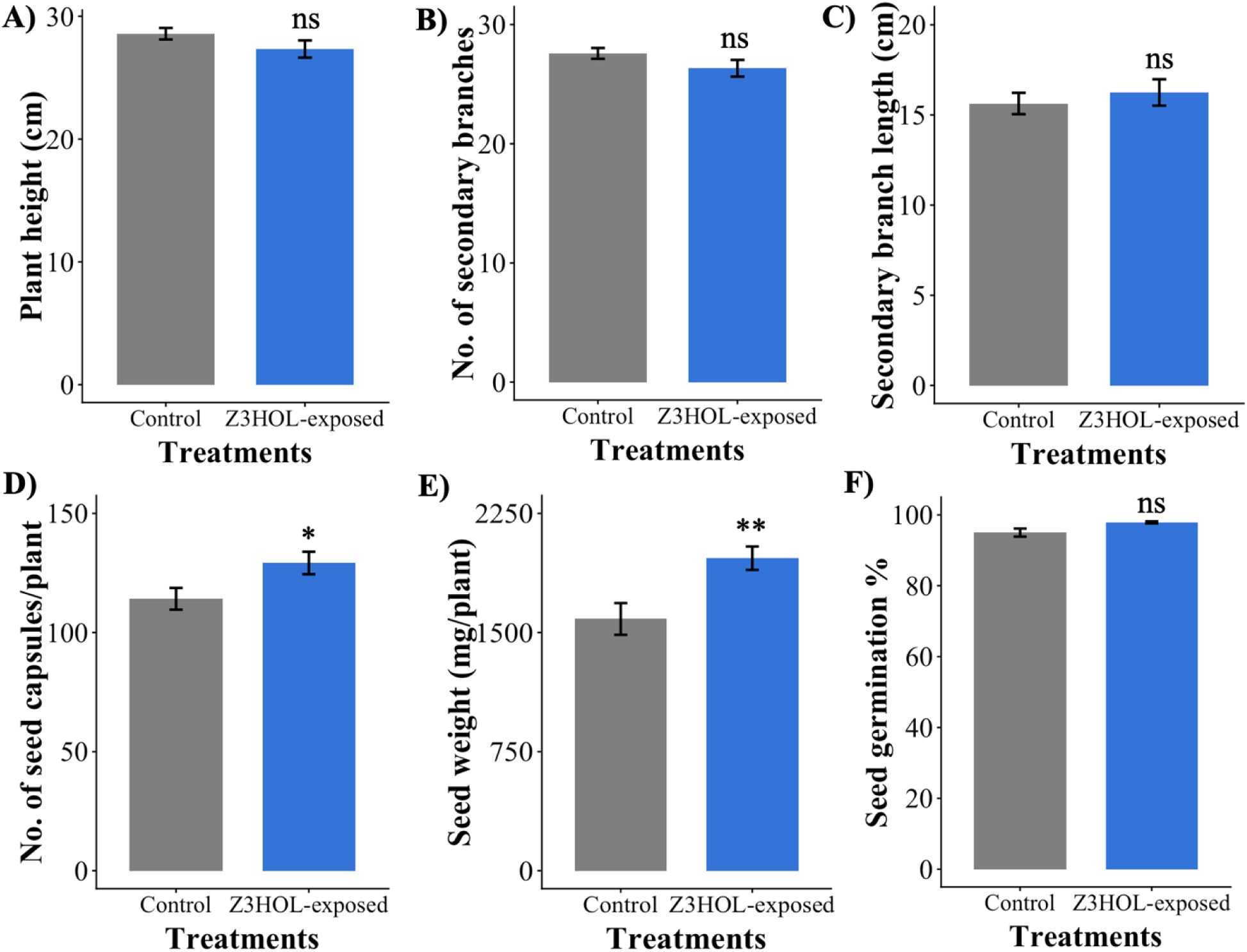
Effect of Z3HOL-exposure on *N. benthamiana* growth and development. **A)** Plant height (cm), **B)** Average number of secondary branches per plant, **C)** Average length of secondary branch (cm), **D)** Number of seed capsules per plant, **E)** Seed weight per plant, and **F)** seed germination percentage. The bar represents mean ± SE (n = 12). * and ** indicate significant difference between control and Z3HOL-exposure at *p* < 0.05, *p* < 0.01, respectively, according to Student’s *t* test.

## 4. Discussion

Herbivore induced plant volatiles (HIPV) play an important role in plant defense priming against insect herbivores. To identify the key volatile component required for plant defense priming, most studies used synthetic compounds at high concentrations, which led to controversies for many years (Dicke *et al*. 2003). Using transgenic plants impaired in their ability to emit specific volatile groups provides a robust approach for dissecting the role of individual volatiles in defense priming. In this study, we used *N. benthamiana* plants with silenced Green leaf Volatile (GLV) biosynthetic pathway (i.e, LOX3 gene silenced plants) to study the role of GLV as airborne priming signals. We assessed priming outcomes using insects’ behavioral and physiological responses as indicators of defense activation, and plants growth and reproductive traits to evaluate potential fitness costs or benefits associated with priming.

Our results demonstrate that when GLV production was silenced in emitter plants, receiver plants no longer exhibited enhanced terpene priming and also showed reduced JA accumulation following simulated herbivory. However, exposure to a physiologically realistic concentration of (*Z*)-3-hexenol (Z3HOL) was sufficient to prime receiver plants for increased JA and terpene production following herbivore challenge. Together, these results indicate that GLV are necessary for full terpene priming and contribute substantially to JA amplification, while Z3HOL alone is sufficient to prime these defense responses. We further demonstrated that this priming has clear ecological consequences. *M. sexta* larvae consumed less leaf tissue and exhibited reduced growth on tobacco plants exposed to Z3HOL or the full HIPV blend, confirming that volatile-mediated priming translated to enhanced resistance. Importantly, we observed no evidence of fitness cost associated with priming. Z3HOL-exposed plants showed no differences in vegetative growth compared to air-exposed controls but produced more seed capsules and greater seed mass. This suggests a positive or neutral trade-off between defense priming and reproduction. Consistent with the definition of priming, elevated production of terpenes and JA was observed only after simulated herbivory, rather than immediately following volatile exposure, indicating primed rather than directly induced defense response (van Hulten *et al*. 2006).

*N. benthamiana* plants emit four main components of GLV, such as (*Z*)-3-hexenal, (*E*)-2-hexenal, (*Z*)-3-hexenol, and (*Z*)-3-hexenyl acetate, after *M. sexta* feeding damage. Among these, only exposure to Z3HOL at physiologically relevant concentration (500 ng h^-1^) primed *N. benthamianana* for enhanced defense against herbivory. However, exposure to (*Z*)-3-hexenal, (*E*)-2-hexenal, and (*Z*)-3-hexenyl acetate neither directly induced nor primed plant defenses against herbivory. These results differ from studies using higher concentration of GLV, which reported direct induction of defense gene expression and increased resistance to herbivores (Farag *et al*. 2005; Yamauchi *et al*. 2018). Moreover, moderate concentrations of (*Z*)-3-hexenyl acetate have been shown to prime *Zea mays* (maize) and *Populus deltoides × nigra* (hybrid poplar) for enhanced JA production and JA-responsive defense genes expression after herbivory (Engelberth *et al*. 2007; Frost *et al*. 2008). This suggests that plants may have evolved species-species mechanisms for responding to volatile cues.

Although Z3HOL alone was sufficient to prime *N. benthamiana*, exposure to the full blend of HIPV consistently primed for stronger defense responses than the exposure to Z3HOL alone. This pattern also aligns with LOX2-silencing results, where removal of the GLV fraction from the complex HIPV blend reduced the magnitude of JA priming but did not abolish it, indicating that additional volatile may have partially compensate for the loss of GLV. Previous studies observed the similar pattern, where exposure to indole and (*Z*)-3-hexenyl acetate induced stronger defense in *Z. mays* than single compounds (Hu *et al*. 2019). These findings support the concept that the full blend of HILV provides a more reliable indicator of herbivore presence than individual volatiles, as proposed by Douma *et al*. (2019).

Behavioral assays further revealed that *M. sexta* preferred to feed on air-exposed plants over Z3HOL- or HIPV-exposed plants. While *M. sexta* larvae initially showed no discrimination among treatments, after several hours, they settled on the leaves from the control plant over Z3HOL- or HIPV-exposed plants. We expected that, after herbivore attack, Z3HOL- or HIPV-exposed plants emitted volatiles in larger amounts than the control; and therefore, be less preferred by *M. sexta* larvae. HIPV-exposed plants do not only accelerate the production of volatiles after herbivore damage but also absorb and reemit significantly higher amounts of volatiles than air-exposed plants for a certain amount of time without herbivore damage (Paudel Timilsena *et al*. 2020). Yet, the higher emission of the volatiles from HIPV-exposed plants did not affect *M. sexta* settlement at the beginning of the experiment. This suggests that feeding preference of *M. sexta* is based on their gustatory response rather than olfactory response alone. The *M. sexta* larval olfactory response to volatiles in our study is in line with previous studies showing that HIPV induced by lepidopteral herbivore do not repel conspecific larvae (Carroll *et al*. 2006, 2008; von Mérey *et al*. 2013). Although not repelled by volatiles, *M. sexta* larvae consumed significantly less leaf materials from Z3HOL- or HIPV-exposed plants than from unexposed control plants, suggesting induction of feeding deterrents or toxic compounds. Reduced herbivore feeding on volatile-exposed plants has been widely reported under both laboratory and field conditions. Under laboratory conditions, Arimura *et al*. (2000) reported less feeding damage by *Teranychus urticae* on *Phaseolus lunatus* plants that were previously exposed to herbivore-damaged conspecific plants. Under field conditions, *N. attenuata* plants and sagebrush suffered less damage from naturally occurring herbivores when exposed to volatiles from artificially damaged sagebrush plants (Karban *et al*. 2000; Karban *et al*. 2006). Similarly, *Salix exigua* and *S. lemmonii* exposed to volatiles from mechanically damaged conspecific plants also suffered less herbivore damage in the fields (Pearse *et al*. 2013). The reduced feeding preference by herbivores may be mediated by direct toxicity of terpenes or presence of antifeedant compound/toxin or both, but this would require additional data to demonstrate.

Consistent with reduced feeding, *M. sexta* larvae grew poorly on plants exposed to Z3HOL or HIPV. Reduced herbivore performance on the primed/volatile-exposed plants has been reported across multiple systems, including reduced growth of *Spodoptera littoralis* on maize (von Mérey *et al*. 2013) and reduced consumption and weight gain of *Plagiodera versicolora* on willow (Yoneya *et al*. 2014). Interestingly, such reductions in herbivore performance can be achieved by exposing herbivores to mixture of GLV (von Mérey *et al*. 2013) and treating plants with individual GLV, such as Z3HOL (in this study) and (Z)-3-hexenyl acetate (Hu *et al*. 2019).

One likely explanation for the reduced weight gain of *M. sexta* on volatiles-exposed plants is enhanced activation of plant defenses. Since JA plays a central role in regulating direct defenses against herbivores (Yang *et al*. 2019), we quantified JA levels in volatile-exposed plants. Our results show that the Z3HOL- and HIPV-exposed plants accumulated significantly higher amounts of JA at the wounding sites than the air-exposed control plants. Consistent with our findings, Z3HOL-exposure has been shown to increase the expression of JA-biosynthesis related genes and JA accumulation in several plants, including *Z. mays* (Farag *et al*. 2005; Engelberth *et al*. 2013), *Camellia sinensis* (Xin *et al*. 2016; Chen *et al*. 2020), and *Solanum lycopersicum* (Su *et al*. 2020; Yang *et al*. 2020). Although, we did not directly test how JA regulated specific defense traits in this study, numerous studies showed that JA activates genes encoding antiherbivore proteins and defense metabolites (Engelberth *et al*. 2004; Farag *et al*. 2005; Ton *et al*. 2007; Frost *et al*. 2008). For instance, exogenous Z3HOL treatment primes JA accumulation and induces JA-responsive genes encoding digestion inhibitors such as proteinase inhibitors, polyphenol oxidases and xylanase inhibitors, and enhance resistance against chewing herbivores (Engelberth *et al*. 2013; Xin *et al*. 2016). Altogether, these findings suggest that enhanced JA accumulation in the Z3HOL- and HIPV-exposed plants likely contributed to increased production of anti-herbivore defense metabolites and reduced larval performance.

In addition to enhanced direct defense, Z3HOL- and HIPV-exposed plants emitted higher levels of defensive volatiles than control plants after herbivory, potentially increasing attraction of natural enemies. (*E*)-α-bergamotene and (*E*)-β-farnesene, two major sesquiterpenes emitted by herbivore damaged *N. benthamiana* plants (Paudel Timilsena *et al*. 2020), were strongly induced in HIPV- and Z3HOL-exposed plants after herbivore damages. Previous studies have shown that (*E*)-α-bergamotene reduces herbivore population on *N. attenuate* by attracting a generalist predator and deterring ovipositing *M. sexta* moths (Kessler & Baldwin 2001; Halitschke *et al*. 2008). Similarly, transgenic *A. thaliana* expressing maize terpene synthase (ZmTPS10), gene that produce (*E*)-α-bergamotene and (*E*)-β-farnesene, attracted the parasitoid *Cotesia marginiventris* and reduced performance of *S. littoralis* (Schnee *et al*. 2006). Thus, volatile exposure (Z3HOL and HIPV) likely enhances both direct and indirect defense against herbivores.

Despite enhanced defense activation, Z3HOL exposure did not negatively affect plant growth and development. Instead, Z3HOL-exposed plants produced more seed capsules, which significantly increased seed weight. Similar fitness benefits have been reported in higher seedling survivorship, branch growth and inflorescence production in sagebrush and *N. attenuata* plants primed by volatiles from clipped neighbors, which showed increased branch growth, numbers of flower and seed production (Karban & Maron 2002; Karban *et al*. 2012). In addition, recent evidence shows that GLV exposure can enhance plant growth and yield through jasmonate-dependent plant-soil feedback that alter rhizosphere microbial communities (Hu et al., 2025). These results suggest that defense priming does not necessarily impose growth or reproductive penalties. If priming were inherently costly, primed plants would be expected to show reduced reproductive fitness due to the cost of secondary metabolite production (reviewed in Guo *et al*. 2018). However, primed plants enhance defense metabolite production only after herbivore attack. Under herbivore-free conditions, as in this study, Z3HOL exposure did not directly induce defense, allowing plants to allocate resources to growth and reproduction. Although priming cost may become apparent under stressful conditions, such as soil nutrient deficiency, low light, drought, frost or herbivory (Karban & Maron 2002; Engelberth & Engelberth 2019; Yip *et al*. 2019), priming with GLV (Z3HAC) can temporarily reduce leaf growth and shift plant physiology toward defense (Engelberth & Engelberth 2019), no such costs were detected under the controlled conditions in this study.

Exposure to Z3HOL enhanced reproductive fitness in *N. benthamiana*. Although the reason behind the enhanced reproductive fitness is unclear, we speculated that JA, which calibrates growth-defense balance (Guo *et al*. 2018), might have shifted resource allocation towards reproduction in the absence of herbivory. JA signaling is known to regulate flower development, especially flower abscission and male sterility (Acosta *et al*. 2009; Song *et al*. 2011; Oh *et al*. 2013; Niwa *et al*. 2018). For example, *N. attenuate* plants with impaired JA signaling (NaJAZd-silenced) increased flower abscission in later stages of flower development which resulted in reduced number of seed capsules and seed mass than wild type plant (Oh *et al*. 2013). Similarly, the constitutive level of JA and its conjugate JA-Ile were much higher in flower buds than those in wounded leaves (Hause *et al*. 2000; Suza *et al*. 2010), suggesting importance of JA signaling in plant reproductive development. Here, our data showed that Z3HOL-exposure increased JA accumulation in leaves after herbivore attack. However, when a primed plant is not under attack, plants could shift resources for reproduction. Whether Z3HOL-exposure also promoted JA biosynthesis in flowers and directly affected flower and seed capsule development, remains to be elucidated.

In summary, silencing GLV production in emitter plants abolished priming of JA accumulation and terpenes emission in receiver plants after simulated herbivory. This demonstrates that GLVs are necessary for defense priming in *N. benthamiana* plants under the experimental condition tested. Moreover, exposure to physiologically relevant concentration of Z3HOL, a common GLV, alone was sufficient to enhance herbivore resistance and reproductive fitness in *N. benthamiana.* Since defense was activated only upon herbivore attack, primed plants did not incur detectable fitness cost under herbivore free conditions. Further long-term field studies are needed to determine whether Z3HOL-mediated priming consistently confers benefits under variable environmental biotic stress, and whether it can be effectively exploited for sustainable pest management.

## Acknowledgments

The authors would like to thank the Department of Entomology and the Center for Chemical Ecology at The Pennsylvania State University for providing support to this study. We dedicate this paper to the memory of late Prof. James H. Tumlinson, who supervised this project with dedication until the end.

## Author Contribution

Conceptualization; BPT, ISA, and JHT. Methodology; BPT, ISA, and SRH. Formal Analysis; BPT and ISA. Visualization; BPT. Writing - original draft: BPT. Writing – Review & Editing; BPT, ISA, GF, SRH, and JHT. Supervision; GF and JHT.

## Data Availability

The datasets generated and analyzed during the current study are available from the corresponding author upon reasonable reques.

## Supporting Information

**Fig. S1 Silencing of the phytoene desaturase gene (PDS) using VIGS in *N. benthamiana*.** *N. benthamiana* plants were syringe infiltrated with TRV2 with PDS fragments (TRV::PDS) and photographed 25 d post-infiltration. TRV::PDS inoculated plants had newly emerged leaves with the photo bleached phenotype characteristic of PDS silencing.

**Fig. S2 Slow-releasing dispenser designed to release physiologically relevant amount of GLV. A)** A mixture of (*Z*)-3-hexenol and lanolin in 1:20 ratio (v/v) inside a GC-insert (25 μL of a mixture per insert), **B)** Slow releasing dispenser, **C)** Volatile exposure set up, **D)** Amount of GLVs (mean ± SE, n = 6) released by *M. sexta* damaged *N. benthamiana* plants at 48 h after feeding initiation, **E)** Amount of individual GLV released from the slow-releasing dispensers (mean ± SE, n = 6).

**Fig. S3 Comparison of constitutive volatiles emissions from undamaged wild-type (WT) plants and *M. sexta*-damaged WT plant.** Volatiles emitted by undamaged plants were below the detection limit of GC-FID.

**Fig. S4 Exposure to (*Z*)-3-hexenal (Z3HAL) alone did not primed *N. benthamiana* plants for enhanced production of monoterpene, sesquiterpene and JA.** *N. benthamiana* plants were exposed to volatiles from *M. sexta* damaged WT (green), undamaged WT plants (gray) or synthetic Z3HAL (dark purple) for 48 h. After 48 h of exposure to volatiles, the receiver plants were transferred to individual clean bell jars and challenged repeatedly by mechanical wounding followed by *M. sexta* regurgitant application. The graph shows the total amount of **A)** monoterpenes and **B)** sesquiterpenes emitted from the challenged receiver plants at different time intervals after initial challenge. Values represent mean ± SE (n = 4). Data were analyzed with a mixed model for repeated measures. Different letters indicate significant (*p* < 0.05) differences between treatments within time points, with Bonferroni’s correction for multiple comparisons. Shaded areas indicate nighttime volatile collections. **C)** For JA analysis, after 48 h of exposure, receiver plants were challenged with mechanical damage followed by *M. sexta* regurgitant application. Wounded portions of the leaf were harvested 30 min after challenge and JA accumulation in the harvested tissue was analyzed (ANOVA, Tukey HSD, n = 4, *p* < 0.05).

**Fig. S5 Exposure to (*Z*)-3-hexenyl acetate (Z3HAC) alone did not primed *N. benthamiana* plants for enhanced production of monoterpene, sesquiterpene and JA.** *N. benthamiana* plants were exposed to volatiles from *M. sexta* damaged WT (green), undamaged WT plants (gray) or synthetic Z3HAL (purple) for 48 h. After 48 h of exposure to volatiles, the receiver plants were transferred to individual clean bell jars and challenged repeatedly by mechanical wounding followed by *M. sexta* regurgitant application. The graph shows the total amount of **A)** monoterpenes and **B)** sesquiterpenes emitted from the challenged receiver plants at different time intervals after initial challenge. Values represent mean ± SE (n = 4). Data were analyzed with a mixed model for repeated measures. Different letters indicate significant (*p* < 0.05) differences between treatments within time points, with Bonferroni’s correction for multiple comparisons. Shaded areas indicate nighttime volatile collections. **C)** For JA analysis, after 48 h of exposure, receiver plants were challenged with mechanical damage followed by *M. sexta* regurgitant application. Wounded portions of the leaf were harvested 30 min after challenge and JA accumulation in the harvested tissue was analyzed (ANOVA, Tukey HSD, n = 4, *p* < 0.05).

**Fig. S6 Volatile (Z3HOL and HIPV) exposure primed defenses against *M. sexta* larvae.** *N. benthamiana* plants were exposed for 48 h to volatile from undamaged plant (air-exposed, gray), 500ng of Z3HOl (Z3HOL-exposed, red) or HIPVs from *M. sexta* damaged plants (HIPV-exposed, green). Values represent mean mortality rate of 3^rd^-instar *M. sexta* larvae 48 h after releasing them on receiver plants (ANOVA, F = 0.389, n = 6, *p* = 0.689).

**Table S1** List of primer sequences used in this study.

