## Supplementary_data for "Exposure to (*Z*)-3-hexenol primes tobacco plants for faster and stronger defense without negatively affecting their ability to grow and reproduce"

The following Supporting Information is available for this article:

**Figure S1**: Silencing of the phytoene desaturase gene (PDS) using VIGS in *N. benthamiana*.

**Figure S2**: Slow-releasing dispenser designed to release physiologically relevant amount of GLV.

**Figure S3:** Comparison of constitutive volatiles emissions from undamaged wild-type (WT) plants and *M. sexta*-damaged WT plant.

**Figure S4:** Exposure to (*Z*)-3-hexenal (Z3HAL) alone did not primed *N. benthamiana* plants for enhanced production of monoterpene, sesquiterpene and JA.

**Figure S5:** Exposure to (*Z*)-3-hexenyl acetate (Z3HAC) alone did not primed *N. benthamiana* plants for enhanced production of monoterpene, sesquiterpene and JA.

**Figure S6**: Volatile (Z3HOL and HIPV) exposure primed defenses against *M. sexta* larvae.

**Table S1**: List of primer sequence used in this study.

**Fig. S1 Silencing of the phytoene desaturase gene (PDS) using VIGS in *N. benthamiana*.** *N. benthamiana* plants were syringe infiltrated with TRV2 with PDS fragments (TRV::PDS) and photographed 25 d post-infiltration. TRV::PDS inoculated plants had newly emerged leaves with the photo bleached phenotype characteristic of PDS silencing.

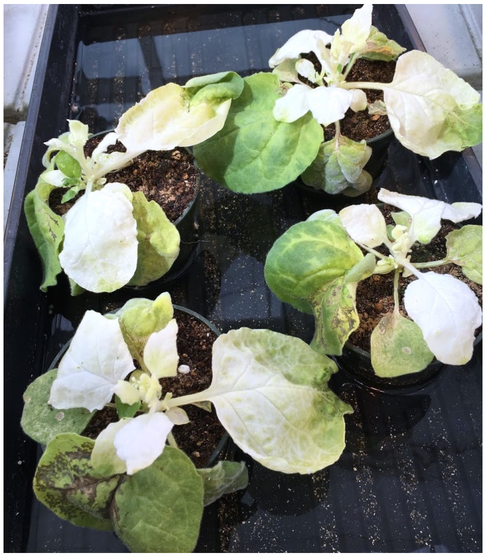

**Fig. S2 Slow-releasing dispenser designed to release physiologically relevant amount of GLV. A)** A mixture of (*Z*)-3-hexenol and lanolin in 1:20 ratio (v/v) inside a GC-insert (25 µL of a mixture per insert), **B)** Slow releasing dispenser, **C)** Volatile exposure set up, **D)** Amount of GLV (mean ± SE, n = 6) released by *M. sexta* damaged *N. benthamiana* plants at 48 h after feeding initiation, **E)** Amount of individual GLV released from the slow-releasing dispensers (mean ± SE, n = 6).

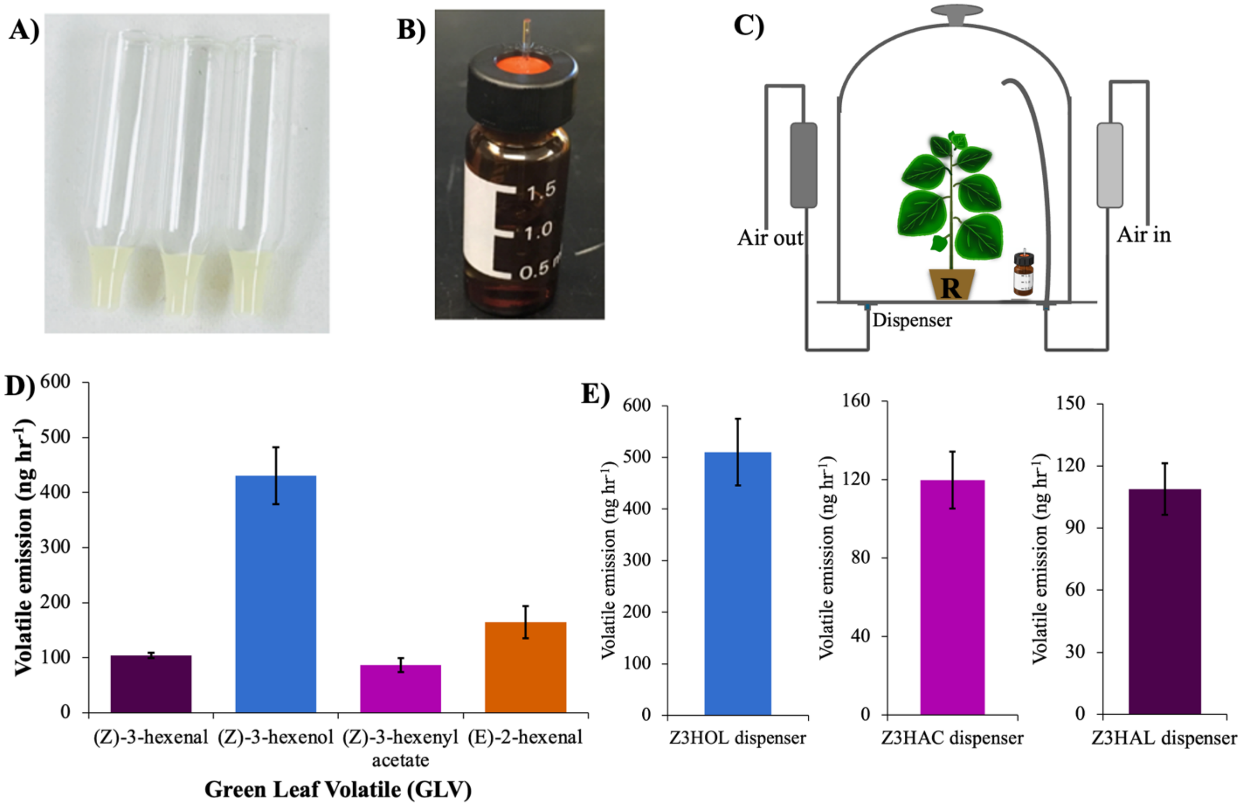

**Fig. S3 Comparison of constitutive volatiles emissions from undamaged wild-type (WT) plants and *M. sexta*-damaged WT plant.** Volatiles emitted by undamaged plants were below the detection limit of GC-FID.

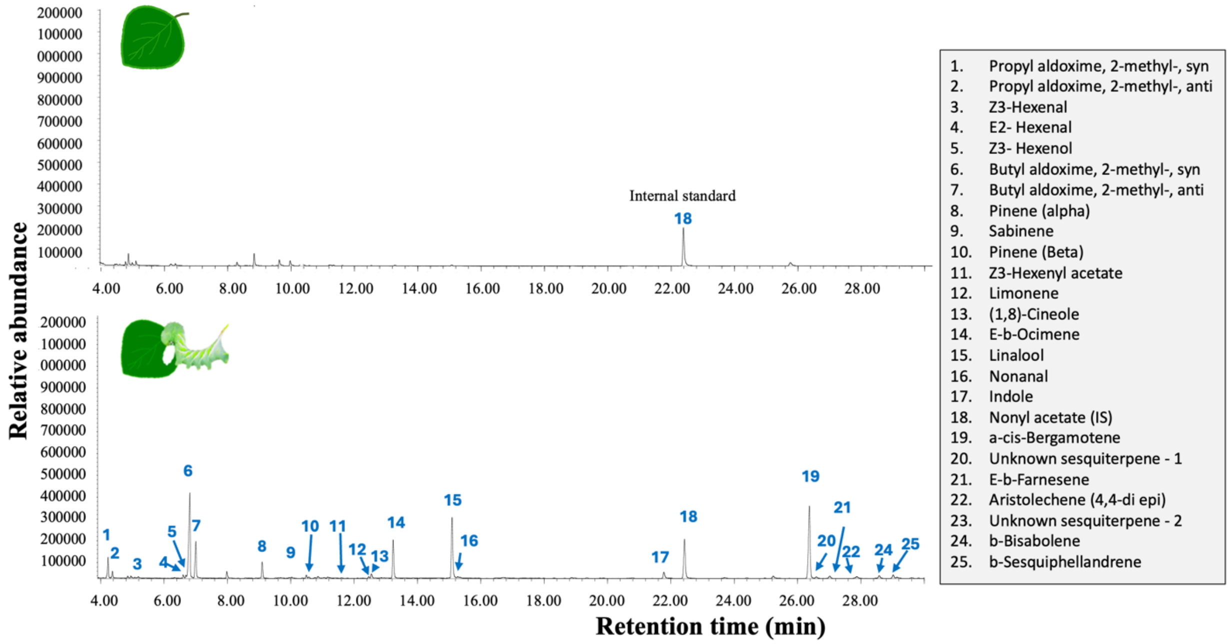

**Fig. S4 Exposure to (*Z*)-3-hexenal (Z3HAL) alone did not primed *N. benthamiana* plants for enhanced production of monoterpene, sesquiterpene and JA.** *N. benthamiana* plants were exposed to volatiles from *M. sexta* damaged WT (green), undamaged WT plants (gray) or synthetic Z3HAL (dark purple) for 48 h. After 48 h of exposure to volatiles, the receiver plants were transferred to individual clean bell jars and challenged repeatedly by mechanical wounding followed by *M. sexta* regurgitant application. The graph shows the total amount of **A)** monoterpenes and **B)** sesquiterpenes emitted from the challenged receiver plants at different time intervals after initial challenge. Values represent mean ± SE (n = 4). Data were analyzed with a mixed model for repeated measures. Different letters indicate significant ($p < 0.05$) differences between treatments within time points, with Bonferroni's correction for multiple comparisons. Shaded areas indicate nighttime volatile collections. **C)** For JA analysis, after 48 h of exposure, receiver plants were challenged with mechanical damage followed by *M. sexta* regurgitant application. Wounded portions of the leaf were harvested 30 min after challenge and JA accumulation in the harvested tissue was analyzed (ANOVA, Tukey HSD, n = 4, $p < 0.05$).

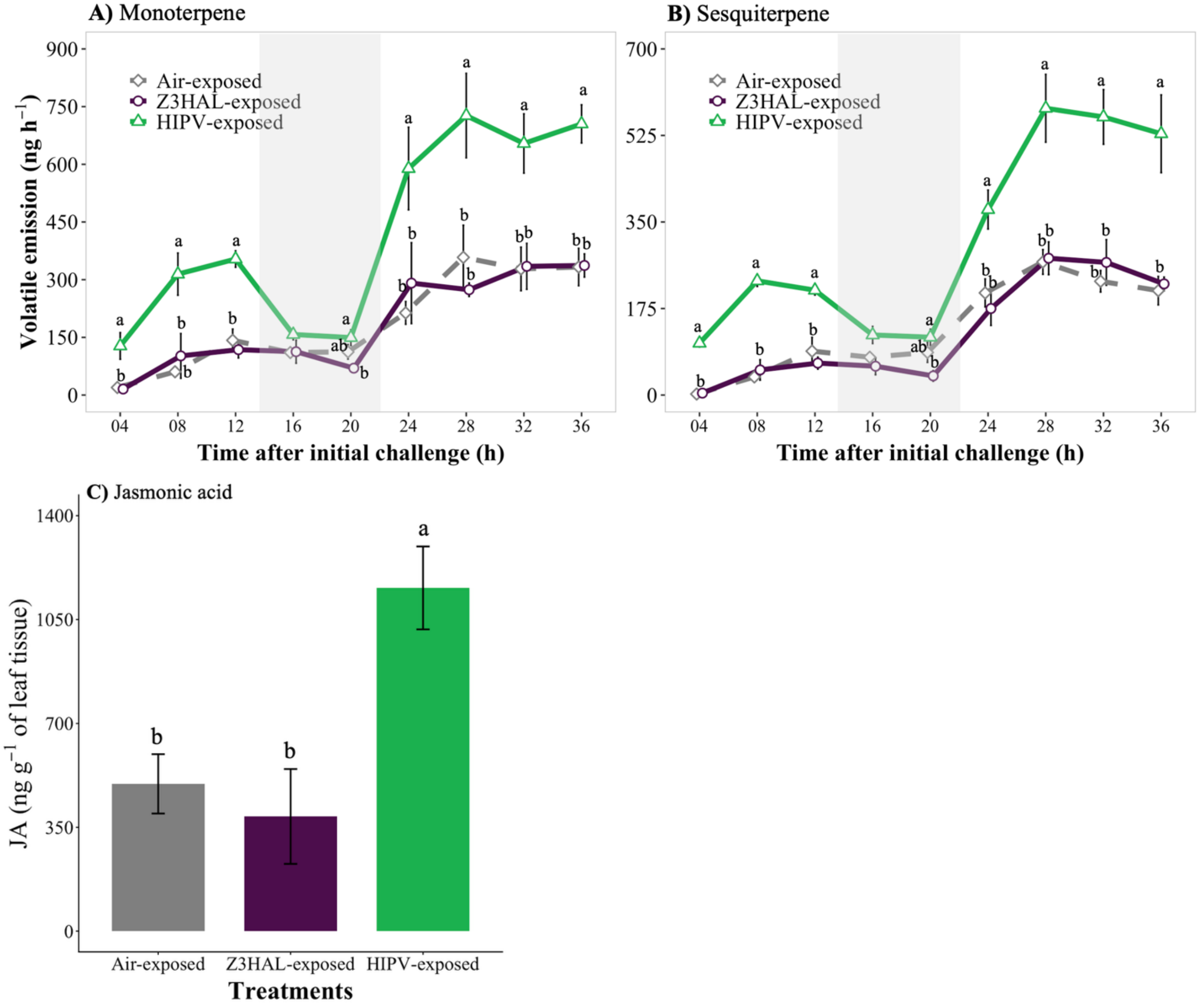

**Fig. S5 Exposure to (*Z*)-3-hexenyl acetate (Z3HAC) alone did not primed *N. benthamiana* plants for enhanced production of monoterpene, sesquiterpene and JA.** *N. benthamiana* plants were exposed to volatiles from *M. sexta* damaged WT (green), undamaged WT plants (gray) or synthetic Z3HAL (purple) for 48 h. After 48 h of exposure to volatiles, the receiver plants were transferred to individual clean bell jars and challenged repeatedly by mechanical wounding followed by *M. sexta* regurgitant application. The graph shows the total amount of **A)** monoterpenes and **B)** sesquiterpenes emitted from the challenged receiver plants at different time intervals after initial challenge. Values represent mean ± SE (n = 4). Data were analyzed with a mixed model for repeated measures. Different letters indicate significant ($p < 0.05$) differences between treatments within time points, with Bonferroni's correction for multiple comparisons. Shaded areas indicate nighttime volatile collections. **C)** For JA analysis, after 48 h of exposure, receiver plants were challenged with mechanical damage followed by *M. sexta* regurgitant application. Wounded portions of the leaf were harvested 30 min after challenge and JA accumulation in the harvested tissue was analyzed (ANOVA, Tukey HSD, n = 4, $p < 0.05$).

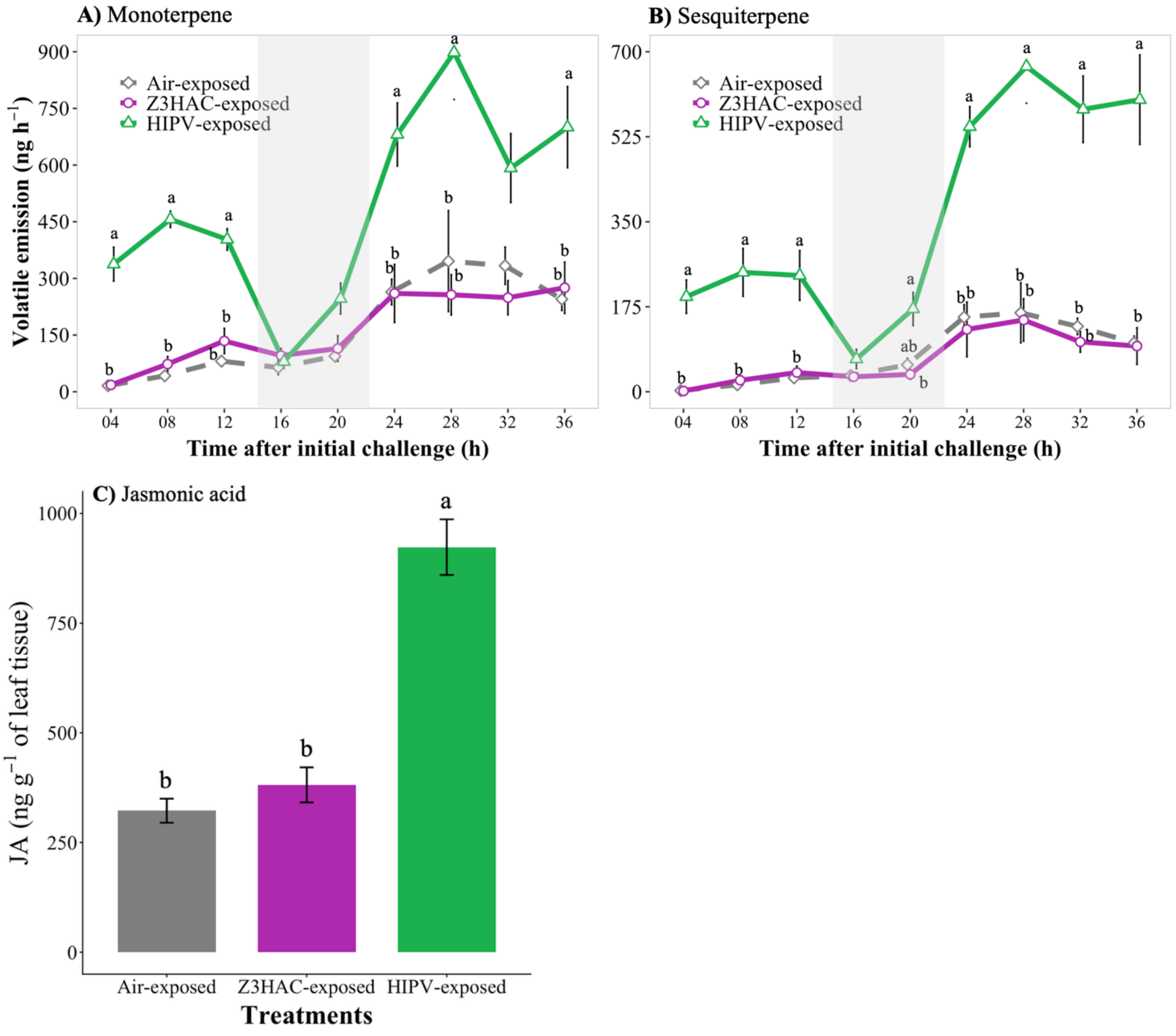

**Fig. S6 Volatile (Z3HOL and HIPV) exposure primed defenses against *M. sexta* larvae.** *N. benthamiana* plants were exposed for 48 h to volatile from undamaged plant (air-exposed, gray), 500ng of Z3HOl (Z3HOL-exposed, red) or HIPVs from *M. sexta* damaged plants (HIPV-exposed, green). Values represent mean mortality rate of 3$^{rd}$-instar *M. sexta* larvae 48 h after releasing them on receiver plants (ANOVA, F = 0.389, n = 6, *p* = 0.689).

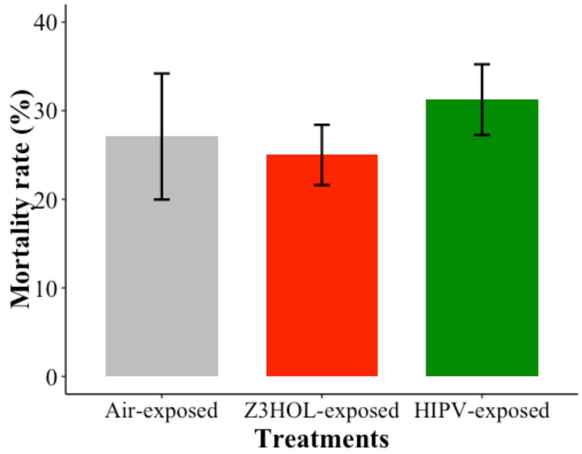


**Table S1: List of primer sequence used in this study.**

| | |
|---|---|
| LOX2 forward | TTTTGGATGTTTTATCAAATCATTCTCCAG |
| LOX2 reverse | GCATCTTATTATGAAATTCATGTCTTGAAAATC |
| LIC vector adaptor forward | CGACGACAAGACCCT |
| LIC vector adaptor reverse | GAGGAGAAGAGCCCT |
